# Opioid-driven disruption of the septal complex reveals a role for neurotensin-expressing neurons in withdrawal

**DOI:** 10.1101/2024.01.15.575766

**Authors:** Rhiana C. Simon, Weston T. Fleming, Pranav Senthilkumar, Brandy A. Briones, Kentaro K. Ishii, Madelyn M. Hjort, Madison M. Martin, Koichi Hashikawa, Andrea D. Sanders, Sam A. Golden, Garret D. Stuber

## Abstract

Because opioid withdrawal is an intensely aversive experience, persons with opioid use disorder (OUD) often relapse to avoid it. The lateral septum (LS) is a forebrain structure that is important in aversion processing, and previous studies have linked the lateral septum (LS) to substance use disorders. It is unclear, however, which precise LS cell types might contribute to the maladaptive state of withdrawal. To address this, we used single-nucleus RNA-sequencing to interrogate cell type specific gene expression changes induced by chronic morphine and withdrawal. We discovered that morphine globally disrupted the transcriptional profile of LS cell types, but Neurotensin-expressing neurons (*Nts*; LS-*Nts* neurons) were selectively activated by naloxone. Using two-photon calcium imaging and *ex vivo* electrophysiology, we next demonstrate that LS-*Nts* neurons receive enhanced glutamatergic drive in morphine-dependent mice and remain hyperactivated during opioid withdrawal. Finally, we showed that activating and silencing LS-*Nts* neurons during opioid withdrawal regulates pain coping behaviors and sociability. Together, these results suggest that LS-*Nts* neurons are a key neural substrate involved in opioid withdrawal and establish the LS as a crucial regulator of adaptive behaviors, specifically pertaining to OUD.

## Introduction

Withdrawal from opioids is intense and highly aversive, which promotes further drug seeking in persons with opioid use disorder (OUD). Anxiety, depression, and increased pain sensitivity are associated with opioid withdrawal, and this collective dysphoric state significantly contributes to relapse^1,2^. Throughout chronic opioid exposure, brain regions that normally support responses to natural rewards and threats undergo widespread neuroadaptations, ultimately shaping the brain in a way that perpetuates opioid misuse^3^. The lateral septum (LS) is a limbic forebrain structure that promotes reinforcement when electrically stimulated^4^ and intense aversion when ablated^5,6^. That is, the LS gates both positive and negative motivation^7^ and is thought to be a key structure involved in substance use disorders^8–13^. Indeed, rodents will self-administer opioids directly into the LS, which suggests that there is a subset of opioid-sensitive neurons in the LS that modulate positive reinforcement^14–16^. However, it remains unclear whether the LS confers withdrawal-induced dysphoria, despite its evident role in fear and stress processing^7^.

Although the LS is primarily a GABAergic hub, it is composed of neuronal subtypes expressing various neuropeptides, receptors and transcription factors that contribute to cell type definition and function^7,17,18^. Activating *Crhr2*-expressing LS neurons is aversive^19^, while manipulating other putative LS subtypes does not seem to evoke the same aversion response^20,21^, further indicating that specific LS cell populations are specialized in processing positive and negative valence stimuli. Chronic opioids also impinge on both reward and aversion circuitry^22^, so uncovering distinct cell types in the LS is imperative for understanding its precise involvement in opioid dependence and withdrawal. While recent studies have targeted various neuropeptide- and receptor-expressing neurons, there is little consensus on LS cell types defined by their transcriptional heterogeneity. To this end, we used single nucleus RNA-sequencing (snRNAseq) and developed a functional atlas to reveal which discrete LS cell populations are recruited during opioid withdrawal. Chronic morphine produced global changes in genes underlying synaptic transmission and protein synthesis across the septum. In contrast, naloxone-precipitated withdrawal most robustly activated *Nts*-expressing LS neurons (LS-*Nts*). Further physiological characterization using two-photon imaging and *ex vivo* electrophysiology revealed that LS-*Nts* neurons remain hyperactivated during prolonged, spontaneous withdrawal. Finally, we demonstrated that bidirectional modulation of LS-*Nts* activity alters opioid withdrawal-induced pain sensitivity and social investigation.

## Results

### Single-nucleus RNA-sequencing reveals a transcriptional atlas for cell types within the septal complex

To shed light on which septal cell types are preferentially disrupted during opioid dependence and withdrawal, we first identified discrete cell types in the septal complex defined by their single cell gene expression. We collected septal tissue from ∼8-week-old male and female C57BL/6J mice and isolated nuclei to generate snRNAseq libraries (**Figure 1A**, **S1A**). In brief, we computationally removed doublets^23^ and corrected for batch effects by utilizing an algorithm based on canonical correlation analysis (CCA)^24^ and mutual nearest neighbors analysis^25^. Data dimensionality was reduced using principal component analysis (PCA) based on the top 3,000 variable genes, which subsequently underwent graph-based clustering (Jaccard-Louvain) and visualized using Uniform Manifold Approximation and Projection (UMAP). Of 53,188 total cells, 32,783 cells were retained as putative neurons based on the expression of *Stmn2* or *Thy1*, canonical neuronal markers typically used to subset neurons^26–28^ (**Figure 1B**, **S1B-C**). After removing potential contaminants and low-quality clusters, we retrieved a final total of 25,607 neurons. Neurons possessed a median unique molecular identifier (UMI) count of 6,075/cell and median gene count of 2,757/cell, consistent with our previously published whole-cell transcriptomic datasets^29–31^ (**Figure S1D-F**). Overall, our snRNAseq data included high quality neurons.

**Figure 1.**
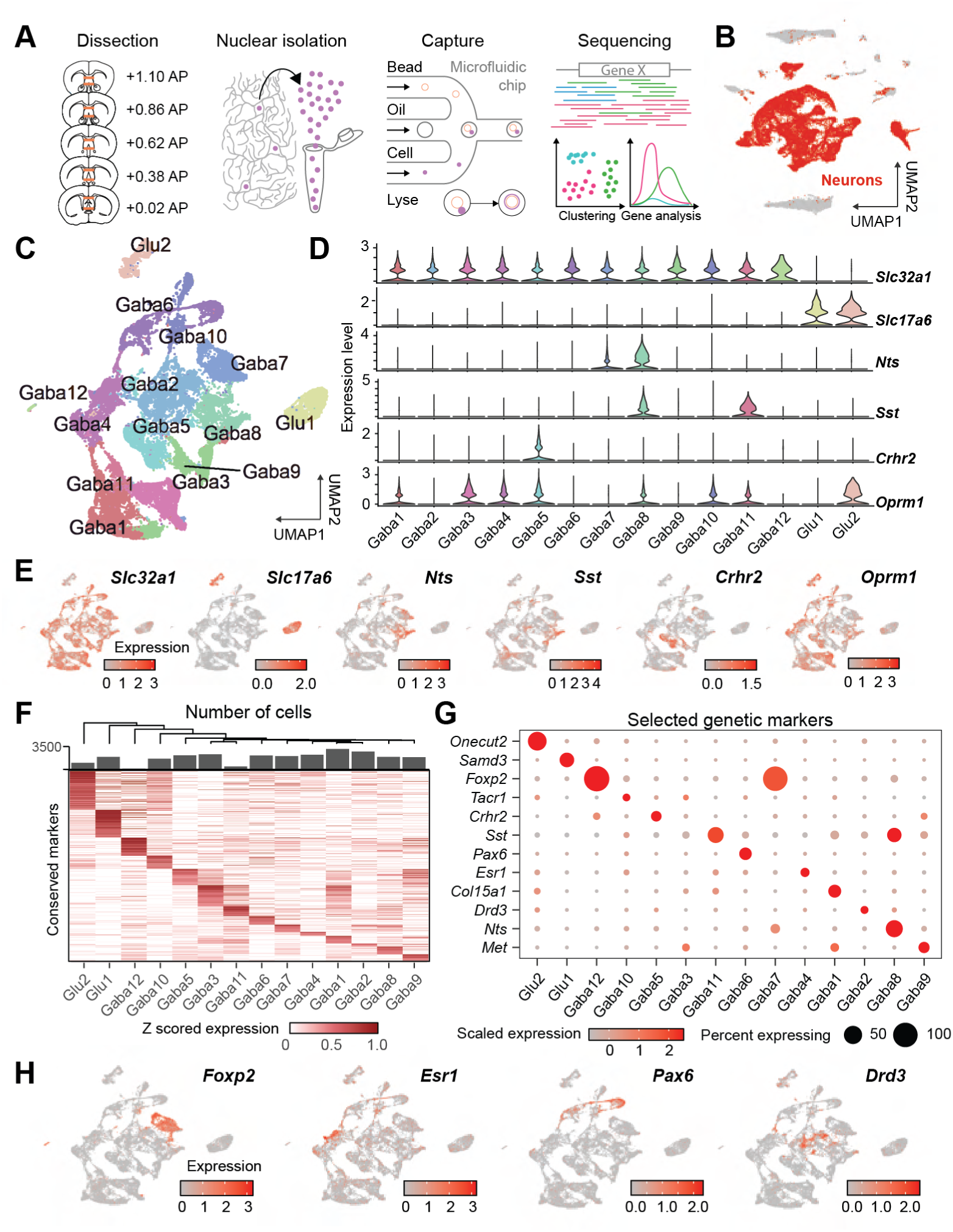
Single-nucleus RNA-sequencing reveals both canonical and undescribed cell populations within the septal complex. (A) Experimental schematic. The septal complex was isolated from male and female C57BL/6J mice. Single nucleus RNA sequencing (snRNAseq) libraries were prepared from extracted septal tissue and sequenced. (B) UMAP plot illustrating individual nuclei clustered according to their transcriptional similarities. Putative neurons are highlighted in red. N = 53,188 nuclei. (C) UMAP plot of individual neurons clustered according to transcriptional similarities. n = 25,607 neurons across 14 bioinformatically-defined clusters. (D) Violin plots of the expression of established LS neuron markers. (E) Feature plots indicating the expression of typical LS neuron markers across space. (F) Heatmap of genes identified to be conserved molecular markers for each neuronal cell type. (G) Discplot illustrating the prevalence of selected candidate markers for each cell cluster. (F) Feature plots indicating the expression of selected candidate markers for Gaba7/12 (*Foxp2+*), Gaba4 (*Esr1+*), Gaba6 (*Pax6+*), and Gaba2 (*Drd3*+).

We identified 14 molecularly defined cell types in the septum (2 glutamatergic and 12 GABAergic), which we determined by evaluating cluster stability over multiple clustering granularities (**Figure 1C**, **S1G**). Among these neurons, we found clusters that corresponded to previously-identified LS cell types, including neurotensin- (*Nts*+; Gaba8), somatostatin- (*Sst*+; Gaba11), corticotropin-releasing hormone receptor 2- (*Crhr2*+; Gaba5) and dopamine receptor 3-expressing (*Drd3*+) neurons^19–21,32–35^. *Crhr2* expression was limited to the Gaba5 subcluster, but *Nts* and *Sst* expression were more promiscuous. Although there was overlap between *Nts* and *Sst* expression (36.4% of *Nts*+ neurons co-express *Sst*), their respective major clusters were transcriptionally divergent (Gaba8 versus Gaba11; **Figure 1F**). Mu opioid receptor (*Oprm1*) was also variably expressed throughout the septum (**Figure 1D-E**), indicating potential sensitivity to both opiates and endogenous opioids^16^. Glu2 was particularly enriched for *Oprm1*, as were Gaba8, Gaba11, and Gaba5 neurons. We next performed conserved marker analysis across all expressed genes to identify gene sets that distinguish each cluster from one another (**Figure 1F**). We chose candidate molecular markers for each cluster based on the relative abundance and expression levels of each conserved gene (**Figure 1G**). Using this approach, we identified novel or understudied neuronal populations within the LS, including those expressing *Foxp2* (Gaba7/12), *Esr1* (Gaba4), *Met* (Gaba9), and *Pax6* (Gaba6).

### Chronic morphine selectively disrupts the transcriptional landscape of the septum

Our snRNAseq data contained experimental groups that we integrated into the same dataset (**Figure 2A**). To model opioid dependence, we used an escalating morphine dose paradigm because this approach has been shown to elicit robust, widespread physiological alterations that underlie drug seeking behaviors^36–38^. An escalating dose of morphine was given to WT C57BL/6J mice for 7 days (10 mg/kg through 70 mg/kg), and on the final day, mice were sacrificed 1 hour after their final dose (Mor group). Control animals were injected with saline (Sal) instead for 7 days and sacrificed on the same timeline. Some Mor subjects were given an injection of naloxone (1 mg/kg) 1 hour following their final morphine injection to precipitate rapid withdrawal and were sacrificed an additional hour later (Nal group). This precipitated withdrawal paradigm produces behavioral phenotypes qualitatively consistent with spontaneous withdrawal during the prolonged absence of opioids^3,37^. Naloxone also standardizes withdrawal responses, whereas spontaneous withdrawal may produce phenotypic variability that contributes to variability in snRNAseq data. After data integration, experimental groups were equally represented across all cell types, and UMI and gene metrics were consistent across groups (**Figure S2A-D**). We also added two additional experimental groups, but we were unable to reconcile sample gene expression differences because these samples were processed using different reagents and sequencing platform (**Figure S2E-F**). However, we maintained these two groups in the base dataset for structural characterization only. See Methods for elaboration.

**Figure 2.**
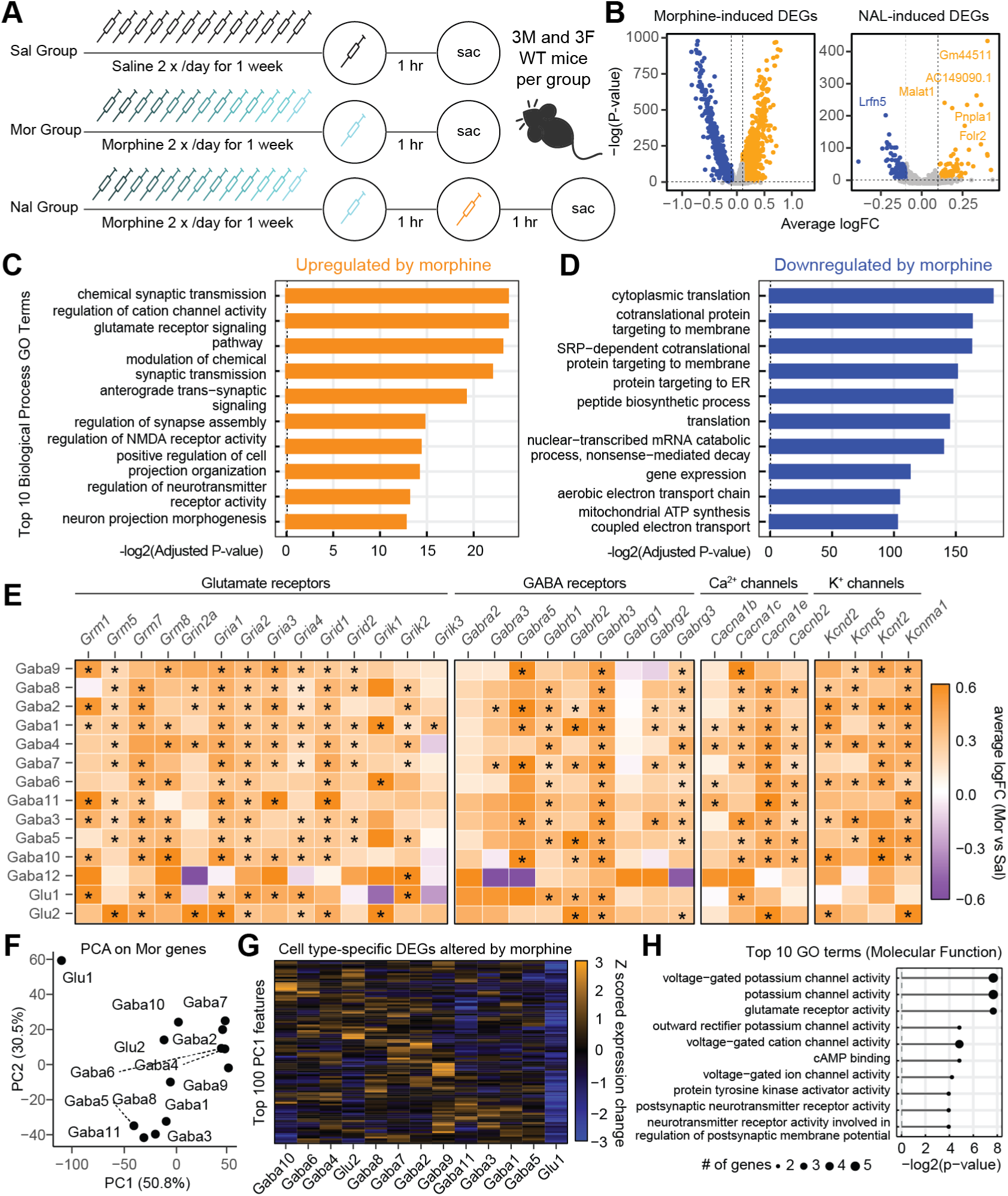
Chronic morphine selectively disrupts the transcriptional landscape of the septum. (A) Experimental schematic. Both male and female C57BL/6J mice were injected with an escalating dose of morphine over the course of 7 days (Mor). Control mice were injected with saline (Sal). To model opioid withdrawal, some animals were injected with 1 mg/kg of naloxone (Nal). (B) Volcano plots indicating the enrichment of DEGs induced by morphine (left, SAL vs MOR) and by naloxone (right, MOR vs NAL) across all cell clusters. P-value cutoff is p < 0.05. DEGs identified by Wilcoxon Rank Sum Test followed by Bonferroni’s post hoc comparison correction. (C) Top 10 Gene Ontology (GO) terms (Biological process) of genes upregulated by chronic morphine. (D) Top 10 GO terms (Biological process) of genes downregulated by chronic morphine. (E) Heatmap illustrating the average logFC between Sal and Mor for different gene classes. “*” indicates significance (Wilcoxon Rank Sum Test followed by Benjamini-Hochberg post hoc comparison correction), and a positive logFC value indicates that the gene is elevated in Mor. (F) Principal component analysis of the logFC expression of DEGs between the Sal and Mor groups. (G) Heatmap of the top 100 features identified in PC1 across all cell types. (H) Top 10 GO terms (Molecular function) identified from the top 100 features.

To identify widespread molecular changes induced by chronic opioids, we first ignored cell cluster labels and combined neurons from all clusters into single pools—one for Sal, one for Mor, and one for Nal. We then performed pairwise comparisons between Sal vs. Mor and Mor vs. Nal to reveal differentially expressed genes (DEGs). Chronic morphine induced and repressed hundreds of DEGs in the septum (2,728 total). In comparison, Mor compared to Nal altered only 487 DEGs (**Figure 2B**). Processes involved in synaptic transmission (GO term: 0007268), NMDA receptor activity (GO term: 2000310), and regulation of neurotransmitter receptor activity (GO term: 0099601) were upregulated (**Figure 2C**), whereas genes involved in translation (GO term: 002181 & 006412), protein targeting to the endoplasmic reticulum (GO term: 0045047), and gene expression (GO term: 0010467) were downregulated by chronic morphine (**Figure 2D**). Mor-induced genes included glutamate receptors, GABA receptors, calcium channels, and potassium channels (**Figure 2E**), which were selectively upregulated across many septal cell types. To examine this more closely, we next performed PCA on the DEGs statistically altered by chronic morphine and determined which features (genes) contributed most to cell type variability in DEG expression (**Figure 2F-G**). GO analysis on the top 100 features once again indicated that genes involved in potassium channel activity and glutamate receptor activity (GO terms: 0005249, 0005267, 0008066) (**Figure 2H**) were potentially driving cell type specific responses to a chronic opioid challenge. Together, our findings suggested that chronic morphine treatment may selectively alter synaptic transmission and membrane properties for different septal cell types, which may consequently impact how these cell types respond to opioid withdrawal.

### *Nts*-expressing neurons in the septum are activated during naloxone-precipitated withdrawal

In order to determine which specific LS cell types were modulated by morphine withdrawal, we next examined immediate early gene (IEG) expression, which is a molecular proxy for neuronal activity^39–41^. *Fos* was relatively depleted in our data with expression in only 3% of neurons, possibly due to *Fos* depletion in the nucleus. However, we detected other IEGs, such as *Arc* (5% of neurons), *Jund* (8%) and *Homer1* (47%), to a higher degree than *Fos* (**Figure 3A**), so we next assessed a literature-based IEG inventory^27^ across all our clusters. We first evaluated naloxone-induced expression changes in “canonical” IEGs, such as *Fos*, *Arc*, *Egr1*, etc., by comparing Mor and Nal group-derived cells. Although there was slight IEG induction in certain cell clusters (e.g. Gaba4), we found that none of these expression differences were statistically significant; we instead found other putative stimulus response genes that were significantly up- or downregulated (**Figure 3B**). Gaba8 (*Nts*+) neurons demonstrated the highest IEG induction (8 genes; *Gem, Pim1, Nptx2, Noct, Ackr4, Homer1, Rheb, Ncoa7,* and *Mbnl2*), followed by Gaba1, Gaba7, and Gaba2. Because individual genes may not fully capture activation patterns, we next assessed a combined IEG metric as a proxy for neuronal activity. Using a panel of additional activity-regulated genes that were deemed to be significantly altered by Nal in any cell cluster, we took the average logFC difference in IEG expression for every neuronal subtype and revealed that Gaba8 demonstrated the most robust change in the IEG profile (**Figure 3C**). Interestingly, we performed the same calculation for canonical IEGs and discovered that Gaba4 (*Esr1*+) neurons were the most enriched instead (**Figure S3A**).

**Figure 3.**
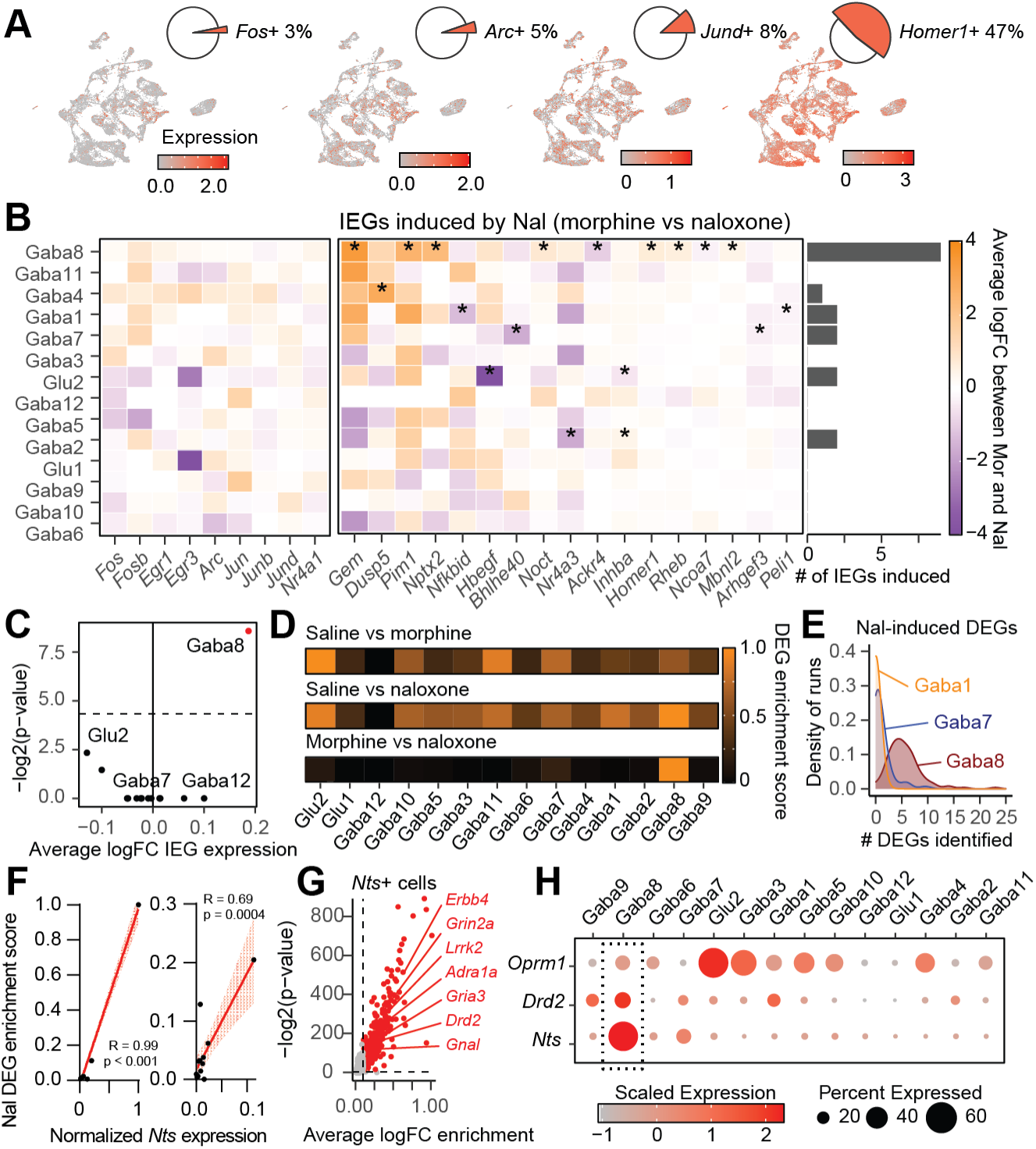
*Nts*-expressing neurons in the septum are recruited during Naloxone-precipitated withdrawal. (A) Feature plots of immediate early genes (IEGs) *Fos*, *Arc* and *Homer1*. Insets, pie charts illustrating the proportion of neurons expressing each IEG (1 copy or greater). (B) Heatmap of IEG induction by Nal across all neuronal clusters. Left panel indicates expression changes of classic IEGs, whereas the middle panel indicates IEGs that are significantly altered in any cell type. Bar graph to the right is the number of IEGs induced within each cluster. Wilcoxon rank sum test followed by Benjamini-Hochberg post-hoc correction. (C) Average logFC expression difference between Mor and Nal of all statistically significantly altered IEGs for each cell cluster. Wilcoxon rank sum test followed by Bonferroni’s post hoc correction. Red indicates statistically-significant clusters. (D) Normalized DEG enrichment scores across cell types for each condition comparison (Sal vs Mor, Sal vs Nal, and Mor vs Nal). (E) Density histogram indicating the frequency of the # of DEGs identified for Gaba7, Gaba8, and Glu2 across 100 runs. (F) The relationship between normalized *Nts* expression (x-axis) and Nal DEG enrichment (y-axis). Left, including Gaba8. Right, excluding Gaba8. (G) Scatterplot indicating the genes that are significantly enriched in *Nts*+ cells versus *Nts*-cells. (H) Discplot of *Oprm1, Drd2, Adra1a,* and *Nts* expression.

Although some IEGs are well-known predictors of neuronal activity, there was variance in IEG expression based on cell cluster. We therefore corroborated our IEG analysis by evaluating naloxone-induced DEG enrichment among each cell cluster, defined by the average number of up- and downregulated DEGs of 100 randomly-sampled cells across 100 iterations. Gaba8 (*Nts*+) possessed the highest DEG enrichment induced by Nal compared to all other cell types (**Figure 3D-E**). To reveal whether cell type clusters were preferentially perturbed across conditions, we next utilized a random forest classifier, an unbiased analysis tool to detect transcriptional changes^42^. AUC scores indicated classifier accuracy, such that a higher score entails a higher degree of transcriptional change. Not only were Gaba8 neurons among the most transcriptionally altered by chronic morphine, but they were also ranked the 2^nd^ most perturbed by naloxone (**Figure S3B-D**). Because the Mor vs. Nal AUC scores were close to 0.50, we also statistically tested for the differences in AUC score distributions to delineate between naloxone-perturbed and naloxone-non-perturbed clusters. Although AUC scores were close to chance, they were significantly elevated in some clusters, notably Gaba8 (*Nts*+), suggesting that the classifier could distinguish between Mor and Nal within specific neuronal subtypes (**Figure S3E**).

Because *Oprm1* was expressed throughout the septum (**Figure 1D**), it was possible that the degree of *Oprm1* expression is linked to septal activation induced by opioids. However, there was no correlation between mean *Oprm1* expression and the Nal-induced DEG score for each cell type (**Figure S3F-G**), suggesting that *Oprm1* RNA expression is not the primary driver for opioid-driven transcriptional alterations in the septum. We instead found a strong, significant relationship between Nal-induced transcriptional changes and *Nts* expression in neuron clusters (**Figure 3F**; with and without Gaba8). Although Gaba8 was the major *Nts* population in our dataset, other cell types such as LS-*Foxp2*, also express *Nts*. We next determined what genes distinguish *Nts*+ from *Nts*-septal neurons, agnostic to cell cluster. Genes involved in chemical synaptic transmission (GO: 0097268) and regulation of neurotransmitter receptor activity (GO: 0099601), specifically genes encoding receptors for catecholaminergic signaling like *Drd2* and *Adra1a*, were enriched in *Nts*+ neurons (**Figure 3G-H**, **S3H**). Given these results, precipitated withdrawal may primarily activate *Nts*+ neurons via inputs from extra-septal regions.

### Discrete LS cell types are organized as a gradient across the medial-lateral and dorsal-ventral axes

Although our snRNAseq data implicated Gaba8 (*Nts*+) neurons as the predominant cell population disrupted during morphine withdrawal, we sought to verify our findings in tissue space. We performed hybridization chain reaction (HCR), which is a highly multiplexed *in situ* hybridization technique that measures many genes simultaneously through sequential rounds of hybridization, amplification, imaging, and probe digestion^43,44^. This approach allowed us to quantify up to 15 different genes within the same tissue. We used septal tissue from 8 naïve WT mice with 15 DNA probes designed to target the following genes: *Nts*, *Drd2*, *Drd3*, *Sst*, *Crhr2*, *Tacr1*, *Foxp2*, *Esr1*, *Met*, *Samd3*, *Col15a1*, *Slc32a1*, *Pax6*, *Onecut2*, and *Slc17a6* (**Figure 4A-B**). Twelve of these genes were molecular markers for snRNAseq-identified cell clusters. Once images were registered onto the same plane, we plotted spatial maps based on the normalized density of cells positive for each gene (**Figure 4C**). Each molecularly defined LS cell type was organized as non-discrete laminae that extend from rostral to caudal aspects of the tissue, which likely arise during the “inside-out” development of the septal complex^45^. *Col15a1*+ and *Onecut2*+ were the densest and most abundant cells, whereas *Esr1*+ and *Pax6+* were the rarest. Incidentally, there was a high degree of overlap among cell populations in the rostral aspects of the LS, but to a lesser degree in caudal regions (**Figure 4D**, **S4A**).

**Figure 4.**
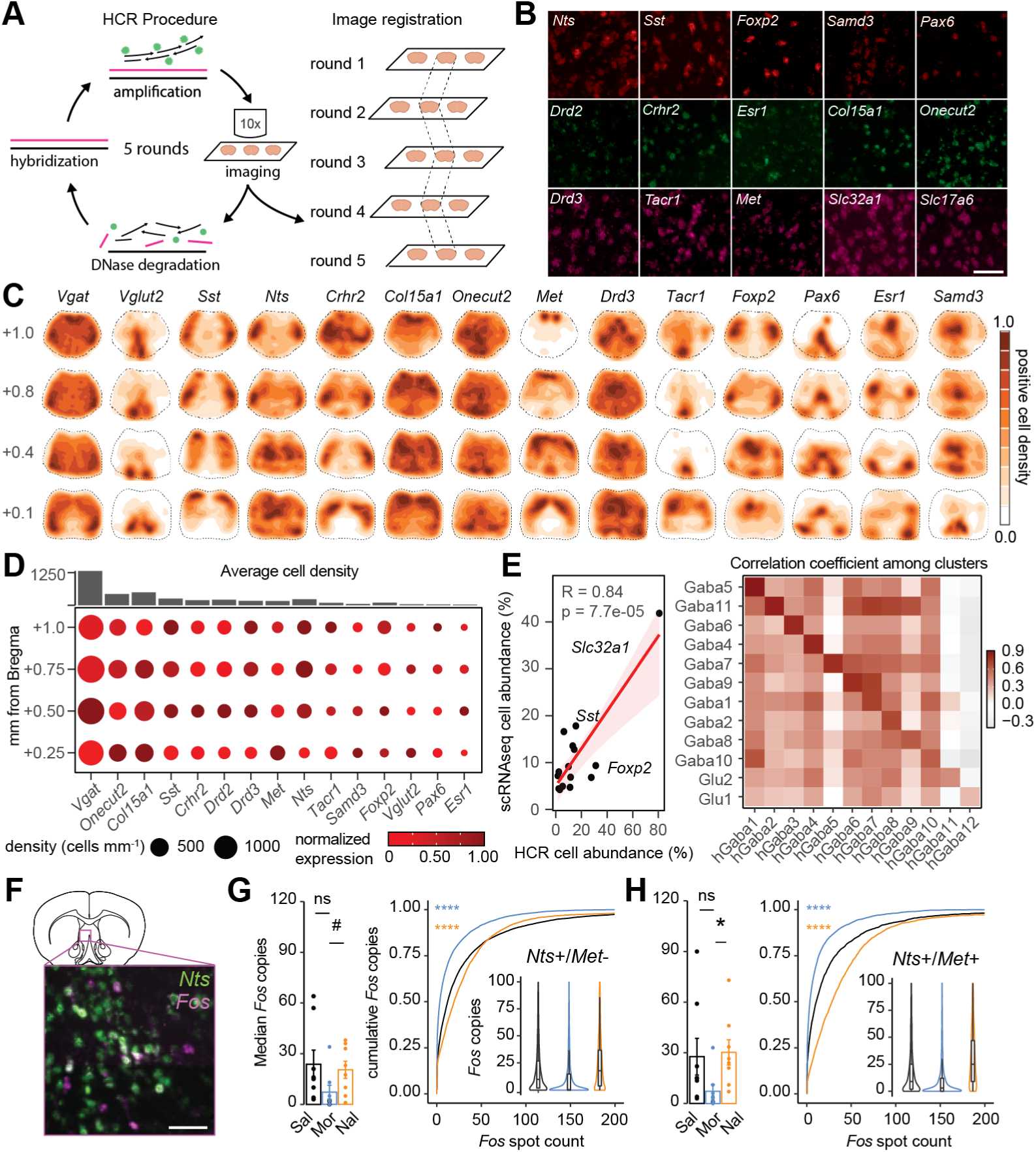
The septal complex possesses gradient-like laminae that may be selectively recruited during opioid withdrawal. (A) Experimental schematic. Septal tissue from 4 male and 4 female C57BL/6J mice were stained for each candidate marker identified from snRNAseq using Hybridization Chain Reaction (HCR). (B) Representative images for each probe organized according to HCR round. Scale bar = 50 μm. (C) Spatial density plots for each gene from all animals collapsed into one dataset. Plots are organized from anterior (∼+1.0 Bregma) to posterior (∼+0.25 Bregma). (D) Disc plot of HCR gene expression across the AP axis of the septum. (E) Left, Correlation between the abundance of cell types identified in snRNAseq versus HCR. Cell type abundance is defined as the percent of cells expressing a given gene (Pearson correlation: R = 0.84. p = 7.7 x 10^-5^). Right, Correlation analysis between clusters defined by snRNAseq (y-axis) and HCR (x-axis). (F) Representative images of *Nts* and *Fos* expression *in situ*. Scale bar = 100 μm. G) Cells positive for *Nts* only. Sal vs Mor: median copies per section (p = 1); median copy difference (p = 1.05 x 10^-10^). Mor vs Nal: median copies per section (p = 0.0897), median copy difference (p = 4.28 x 10^-5^). (H) *Nts*+ and *Met*+ cells. Sal vs Mor: median copies per section (p = 0.551); median copy difference (p = 7.58 x 10^-10^). Mor vs Nal: median copies per section (p = 0.0181); median copy difference (p = 1.71 x 10^-13^). #p < 0.1, *p < 0.05, **p < 0.01, ***p < 0.001, ***p < 0.001, ****p < 0.0001.

To evaluate the congruency between snRNAseq and HCR, we clustered cells based on HCR gene expression data (**Figure S4B-C**). The correlation between the abundance of cells expressing each gene in the snRNAseq and HCR datasets was robust and statistically significant. This correlation was driven primarily by *Slc32a1* expression, but this relationship ultimately extended to a cluster-by-cluster basis as well (**Figure 4E**). Therefore, despite differences in sampling biases of the two techniques, the relative distribution of each cell type was consistent across the two datasets. We next inferred where each snRNAseq cell cluster may anatomically reside within the septal complex. Gaba2 (LS-*Drd3*), Gaba4 (LS-*Esr1*), Gaba5 (LS-*Crhr2*), Gaba7 (LS-*Foxp2*), Gaba8 (LS-*Nts*), and Gaba11 (LS-*Sst*) comprised the LS. Glu1, Gaba6, and Gaba10 represented the medial septum (MS), and Glu2 likely resided in the septofimbrial nucleus of the septum (SFi). Finally, Gaba12 was the smallest cell cluster (**Figure 1F**; n = 112) and was likely a rare cell type within the Septofimbrial nucleus (SFi).

We also verified that naloxone induced IEG expression in LS-*Nts* neurons *in situ*. A new cohort of mice underwent morphine escalation (Mor), morphine withdrawal (Nal), and repeated saline injections (Sal) and were sacrificed 1 hour after injection. We probed for *Fos* expression in both LS-*Nts* and LS-*Met* cells, as both were implicated as naloxone-responsive cell types in our snRNAseq data (**Figure 4F**). While both LS-*Nts* (exclusive of *Met*) and LS-*Met* (exclusive of *Nts*) demonstrated naloxone-induced *Fos* expression, LS neurons expressing both *Nts* and *Met* showed the highest *Fos* induction during precipitated withdrawal (**Figure 4G-H**). Incidentally, cells negative for both were depleted for *Fos* (**Figure S4E-G**). We predicted that if both genes are associated with the largest *Fos* response, then perhaps they were localized to a specific region in the septum. Indeed, we found that neurons co-expressing *Nts* and *Met* resided in the intermediate aspect of the LS (LSi) (**Figure S4H-I**). Altogether, our snRNAseq and HCR analyses revealed LS-*Nts* neurons as robustly activated following naloxone-precipitated withdrawal and are spatially localized to the LSi. While postmortem analyses of brain tissue provided a plethora of molecular and spatial information, these approaches are unable to provide real-time temporal information about how opioid withdrawal may modulate LS-*Nts* activity *in vivo*.

### Opioid withdrawal elevates LS-*Nts* activity *in vivo*

Opioid withdrawal disrupts emotional regulation, producing a multifaceted dysphoric state which includes both anhedonia and hypersensitivity to fear and stress^46,47^. Therefore, we next sought to reveal whether opioid exposure and withdrawal alter LS-*Nts* responsivity to rewarding and aversive stimuli. To this end, we used two-photon imaging in awake, behaving mice, which allowed us to track how single neurons are altered during the emergence of opioid dependence. We injected *Nts*-Cre mice with AAVDJ-CAG-DIO-Gcamp6s unilaterally in the LSi, followed by a GRIN lens implantation directly above the same site (**Figure 5A**). Once mice were habituated to head-fixation, we first imaged mice as drug-naïve to establish baseline calcium dynamics for LS-*Nts* neurons. LS-*Nts* neurons expressing Gcamp6s and were spontaneously active *in vivo* (**Figure 5B**, **S5A-B**). We then exposed mice to interleaved presentations of 10% sucrose (rewarding; 80% of trials) and 20 psi air puffs (mildly aversive; 20% of trials) to evaluate how innately positive and negative valanced stimuli modulate LS-*Nts* activity. Mice were then subsequently split into either saline-treated (Sal) or morphine-treated (Mor) groups. Neuronal activity was recorded every other day during our morphine escalation protocol (**Figure 5C**; D2, D4, D6). Each imaging session occurred before mice received any saline or morphine for that day, as we found that high doses of morphine suppressed appetitive licking for several hours. LS-*Nts* neurons were responsive to both air puffs and sucrose (**Figure 5D**), and Sal and Mor neurons were not significantly different in their responsivity to either stimulus at baseline. While opioid exposure did not influence response magnitude to air puffs and sucrose, it reduced the proportion of cells responding to sucrose and not air puffs on D2 and D4. (**Figure S5C-D**).

**Figure 5.**
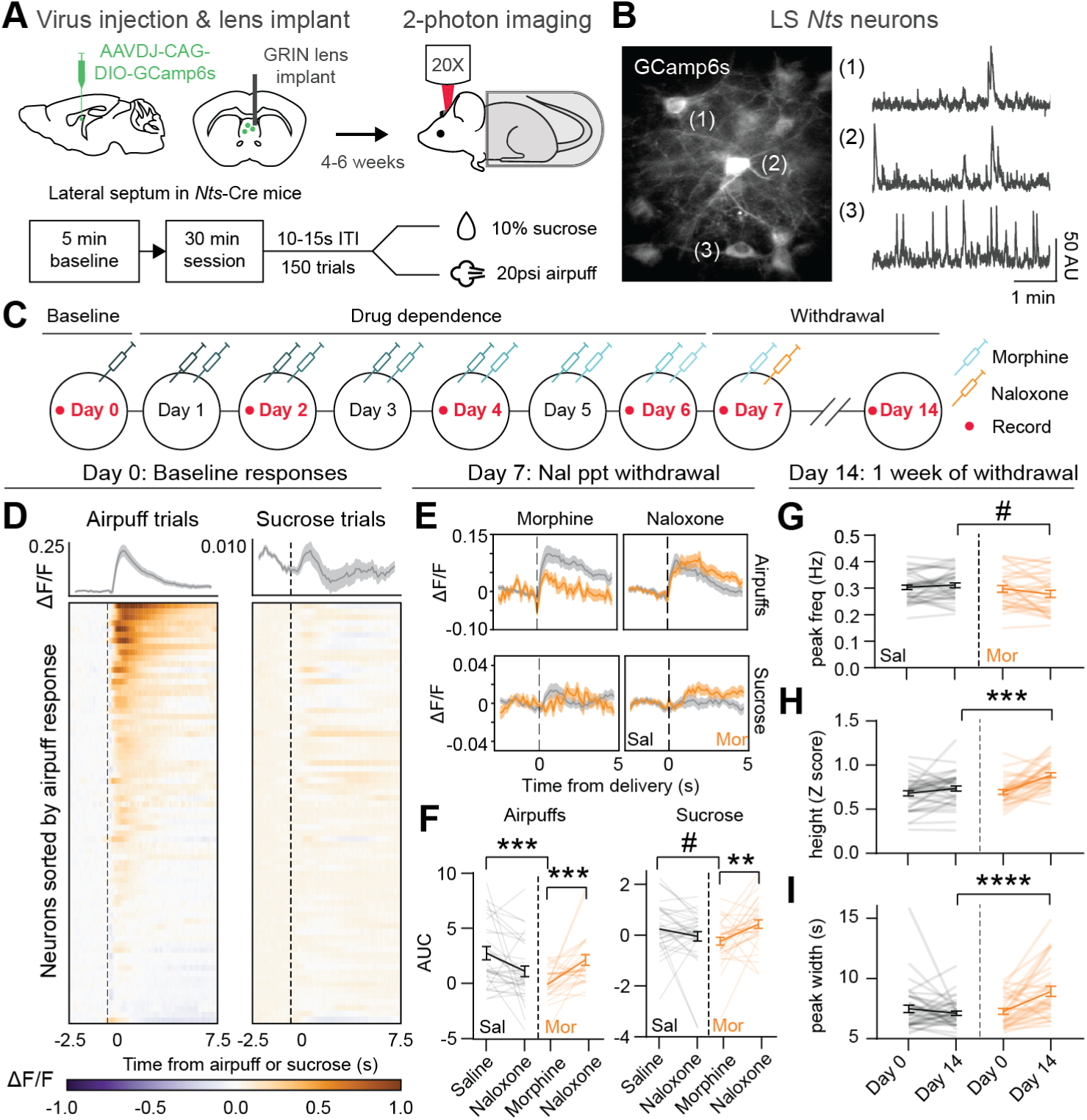
Opioid withdrawal elevates LS-*Nts* activity in vivo. (A) Injection scheme and experimental schematic. 4 male and 2 female *Nts*-cre mice were injected with AAVDJ-hSyn-DIO-GCamp6s and implanted with a 0.5 mm x 7 mm GRIN lens for two-photon imaging. (B) Left, sample mean image of GCamp6s expression in LS-*Nts* neurons from a representative animal. Right, sample raw fluorescence traces of 3 representative neurons. (C) Experimental timeline. Animals were habituated to head-fixation and sucrose consumption for 5-7 days. Baseline data were taken prior to the onset of either saline (Sal) or escalating morphine (Mor) injections. Opioid withdrawal was precipitated by a 1 mg/kg dose of naloxone 1 hour after the final dose of morphine. Sal animals also received naloxone. (D) LS-*Nts* activity evoked by air puffs and sucrose presentations in one representative animal on Day 0. Top, lick rate aligned to air puff or sucrose presentation. Bottom, peri-event time histograms (PSTHs) of neuronal responses to air puffs (left) and sucrose (right). n = 79 cells. (E) Mean naloxone-induced responses of Sal-treated (n = 33) and Mor-treated (n = 32) cells to air puffs (top) and sucrose (bottom). Sal cells received saline followed by naloxone, whereas Mor-treated cells received morphine followed by naloxone. (F) AUC measurements for air puff-evoked (Left) and sucrose-evoked (Right) responses of Sal- and Mor-treated cells before and after naloxone. Left, 2-way RM ANOVA followed by Šídák’s multiple comparisons; Sal vs Mor interaction: F_(1, 63)_ = 22.86, p < 0.0001; Sal vs Mor pre-naloxone, p = 0.002; Mor pre- vs post-naloxone, p = 0.0005. Right, 2-way RM ANOVA followed by Šídák’s multiple comparisons; interaction: F_(1, 63)_ = 10.47, p = 0.0019; Sal vs Mor pre-naloxone, p = 0.0867; Mor pre- vs post-naloxone, p = 0.0047. (G-I) Spontaneous activity analysis of Sal-treated (n = 39) and Mor-treated (n = 32) cells after 7 days of withdrawal (Day 14). (G) Peak frequency. 2-way RM ANOVA followed by Šídák’s multiple comparisons; interaction: F_(1, 69)_ = 4.638, p = 0.0348; Day14 Sal vs Mor, p = 0.0697. (H) Peak amplitude. 2-way RM ANOVA followed by Šídák’s multiple comparisons; interaction: F_(1, 69)_ = 10.51, p = 0.0018; Day14 Sal vs Mor, p = 0.0006. (I) Peak width. 2-way RM ANOVA followed by Šídák’s multiple comparisons; interaction: F_(1, 69)_ = 17.41, p < 0.0001; Day14 Sal vs Mor, p < 0.0001. Summary data are represented as mean ± SEM. #p < 0.1, *p < 0.05, **p < 0.01, ***p < 0.001, ***p < 0.001, ****p < 0.0001.

Our snRNAseq and HCR data suggested that naloxone activates LS-*Nts* neurons. To examine this *in vivo*, we gave mice their final saline or morphine injection on Day7 and recorded LS-*Nts* activity ∼30 min following treatment. Afterwards, Sal and Mor mice were injected with naloxone and re-imaged ∼30 min following naloxone treatment. We found that acute morphine exposure blunted both air puff- and sucrose-evoked responses from LS-*Nts* neurons (**Figure 5E-F)**. Interestingly, naloxone treatment reversed the influence of morphine on LS-*Nts* neurons in opioid-dependent mice, allowing these neurons to recover responses to both sucrose and air puffs (**Figure 5F**). Sal neurons, in contrast, were possibly mildly suppressed following naloxone treatment. While it was evident that Nal rapidly modulated LS-*Nts* activity, it was unclear whether LS-*Nts* activity changes during withdrawal were transient or prolonged. To address this, we re-imaged mice 1 week following the completion of morphine injections. We found that although air puff and sucrose-evoked responses were minimally different between Sal and Mor neurons at 1 week of spontaneous withdrawal (**Figure S5E**), neurons from Mor-treated mice showed higher and longer lasting calcium peaks compared to neurons recorded in Sal-treated mice (**Figure 5G-** I). LS-*Nts* neurons may possess constitutively higher spontaneous activity during sustained abstinence from morphine, indicating LS-*Nts* neurons as a potential regulator of behavioral dysfunction during opioid withdrawal.

### Glutamatergic drive is enhanced at LS-*Nts* neurons during opioid withdrawal

LS-*Nts* activity in morphine-dependent animals was upregulated during spontaneous and protracted withdrawal, an effect that are likely driven by synaptic alterations induced by chronic morphine. Because our snRNAseq data indicated that chronic morphine upregulated GLU receptor gene expression in LS-*Nts* neurons (**Figure 2E**), these neurons may undergo excitatory synaptic strengthening during morphine exposure. In the absence of morphine, LS-*Nts* neurons may then become hyperactivated due to increased glutamatergic drive. Therefore, we next sought to determine whether LS-*Nts* displayed alterations in synaptic transmission using *ex vivo* electrophysiology. To label LS-*Nts* neurons for identification during electrophysiology experiments, we injected AAV5-hSyn-DIO-eYFP bilaterally into the LS of *Nts*-Cre mice (**Figure 6A**). Two weeks later, mice underwent the same morphine escalation paradigm (Mor) or received saline (Sal). All mice received naloxone. 6-10 days later, brain slices were collected for synaptic physiology measurements (**Figure 6A-B**). Mini excitatory and inhibitory postsynaptic currents (mEPSCs and mIPSCs) were recorded in a whole-cell configuration with membrane voltage clamped at -70 mV for mEPSCs and +10 mV for mIPSCs (**Figure 6B**). When possible, both measurements were collected from the same cell. LS-*Nts* neurons from mice that underwent morphine treatment showed increased mEPSC frequency (**Figure 6C-D**) and amplitude (**Figure 6E**) compared to saline-treated mice, suggesting that chronic morphine and withdrawal increases excitatory synaptic strength at these neurons. We also noticed that GABA receptor-related genes were upregulated in LS-*Nts* neurons (**Figure 2E**), but morphine treatment and withdrawal did not affect mIPSC frequency or amplitude at LS-*Nts* neurons (**Figure 6F-H**), suggesting that overall inhibitory synaptic strength at these neurons was unaffected by morphine. Our *ex vivo* electrophysiology experiments demonstrated that opioid dependence and withdrawal enhances glutamatergic signaling onto LS-*Nts* neurons.

**Figure 6.**
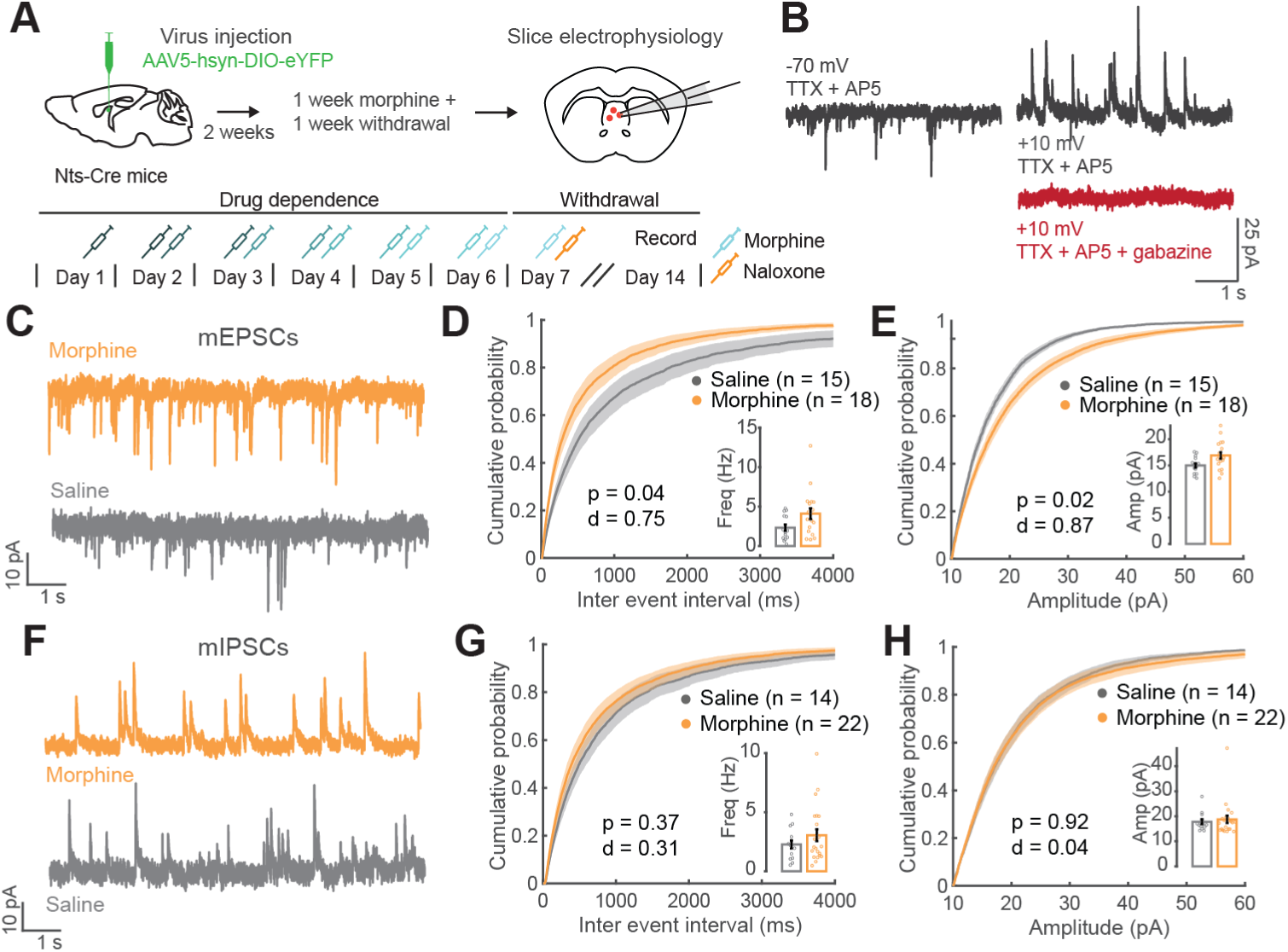
Morphine treatment and withdrawal alter excitatory, but not inhibitory, synaptic strength at LS-*Nts* neurons. (A) Viral schematic for labeling LS-*Nts* neurons and timeline of experiment. (B) Example whole-cell, voltage-clamp traces from an LS-*Nts* neuron. mEPSCs were measured at -70 mV (left) and mIPSCs were measured at +10 mV (right). Currents at +10 mV are abolished by gabazine (bottom). (C) Example traces of mEPSCs. (D) Cumulative probability of inter-event intervals for mEPSCs. Morphine treatment increased mEPSC frequency compared to saline-treated controls (d = 0.749, p = 0.046 for group in an LMER on frequency with group as a fixed effect and mouse and cell as random effects. Inset: median frequency of mEPSCs. Morphine, n = 18 cells, 6 mice, 4.1 Hz ± 0.7. Saline, n = 15 cells, 4 mice, 2.3 Hz ± 0.4). (E) Cumulative probability of amplitudes for mEPSCs. Morphine treatment increased mEPSC amplitude compared to controls (d = 0.866, p = 0.022 for group in an LMER on amplitude with group as a fixed effect and mouse and cell as random effects. Inset: median amplitude of mEPSCs. Morphine, n = 18 cells, 6 mice, 16.9 pA ± 0.6. Saline, n = 15 cells, 4 mice, 15.0 pA ± 0.4). (F) Example traces of mIPSCs. (G) Cumulative probability of inter-event intervals for mIPSCs. Morphine treatment did not affect mIPSC frequency compared to controls (d = 0.310, p = 0.374 for group in an LMER on frequency with group as a fixed effect and mouse and cell as random effects. Inset: median frequency of mIPSCs. Morphine, n = 22 cells, 6 mice, 3.04 Hz ± 0.5. Saline, n = 14 cells, 4 mice, 2.3 Hz ± 0.3). (H) Cumulative probability of amplitudes for mIPSCs. Morphine treatment did not affect mIPSC amplitude compared to controls (d = 0.866, p = 0.022 for group in an LMER on amplitude with group as a fixed effect and mouse and cell as random effects. Inset: median amplitude of mIPSCs. Morphine, n = 22 cells, 6 mice, 18.7 pA ± 1.5. Saline, n = 14 cells, 4 mice, 17.8 pA ± 1.0).

### Silencing LS-*Nts* neurons exacerbates withdrawal-induced pain coping and triggers sexually-dimorphic alterations in sociability

Because we discovered that LS-*Nts* neurons were hyperactive in the absence of opioids, we predicted that elevated LS-*Nts* activity contributes to maladaptive behaviors associated with opioid withdrawal. To address this, we used genetically-encoded tetanus toxin light chain (TetTox) to silence these neurons. TetTox disrupts vesicle release machinery and prevents neurotransmission from treated neurons^48^. Incidentally, TetTox silencing has been functionally validated in another study examining LS-*Nts* neurons in feeding^32^. After injecting 14 animals with AAVDJ-EF1a-DIO-TetTox-GFP (TetTox group) and 12 animals with AAV5-EF1a-DIO-eYFP (control group), we waited 8-10 weeks for viral expression and performed a battery of 6 behavioral tests over a 2-month period (**Figure 7A-B**). Following baseline testing, control and TetTox mice underwent morphine escalation and precipitated withdrawal. We then measured behavioral changes during 1-2 and 4-5 weeks of morphine withdrawal. Using this method, we assessed how chronic loss of LS-*Nts* function influences the emergence of withdrawal phenotypes.

**Figure 7.**
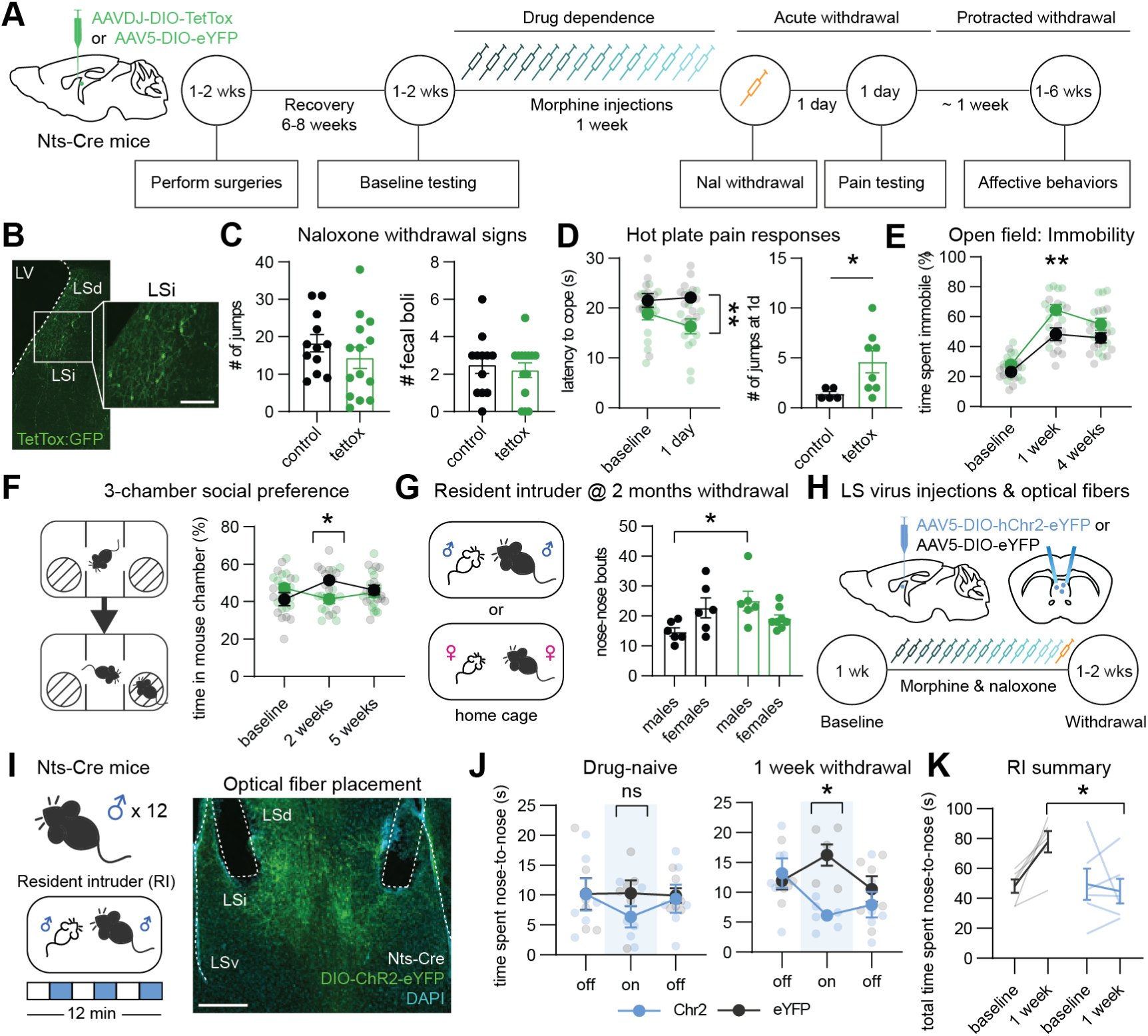
Silencing LS-*Nts* neurons exacerbates withdrawal-induced pain coping and triggers sexually-dimorphic alterations in sociability. (A) Experimental schematic. *Nts*-Cre mice were bilaterally injected with either AAV5-EF1a-DIO-eYFP (control) or AAVDJ-EF1a-DIO-TetTox-GFP (TetTox) into the LS. TetTox was allowed to express for 8 weeks prior to baseline behavior testing, after which all mice underwent 7 days of morphine escalation and naloxone-precipitated withdrawal. (B) Representative image of TetTox-GFP expression in the LSi of a *Nts*-Cre mouse. Scale bar = 200 μm (C) Naloxone-induced withdrawal signs. Left, number of jumps. Unpaired t-test: t_(24)_ = 1.033, p = 0.3118. Right, number of fecal boli. Unpaired t-test: t_(24)_ = 0.4698, p = 0.6427. (D) Hot plate. Left, latency to performing a coping action (hind paw lick or jump). 2-way RM ANOVA followed by Šídák’s multiple comparisons; group effect: F_(1,24)_ = 6.720, p = 0.0160; control vs. TetTox at 1d, p = 0.0069. Right, TetTox increases the number of jumps mice perform at 1 day of withdrawal. Unpaired t-test: t_(11)_ = 2.257, p = 0.0453. (E) Open field mobility. 2-way RM ANOVA followed by Šídák’s multiple comparisons; group effect: F_(1, 24)_ = 13.98, p = 0.0010; control vs. TetTox at 1 week, p < 0.0026. (F) 3-chamber social preference. 2-way RM ANOVA followed by Šídák’s multiple comparisons; interaction: F_(2,48)_ = 5.197, p = 0.0091; control vs. TetTox at 2 weeks, p = 0.0298. (G) Resident intruder assay. Left, nose-to-nose interactions. 2-way RM ANOVA followed by Šídák’s multiple comparisons; interaction: F_(1, 22)_ = 8.433, p = 0.0082; control males vs TetTox males, p = 0.0144. (H) Experimental schematic for optogenetic activation of LS-*Nts* neurons. Male Nts-Cre mice were bilaterally injected with either AAV5-ef1a-DIO-eYFP (control) or AAV5-ef1a-DIO-hChr2-eYFP (stim), and bilateral optical fibers were implanted above the LS. (I) Left, mice underwent the resident intruder task, composed of 2 min long light off and light on epochs, during which 5-7 mW 470 mm laser light was delivered at 20 Hz, 5 ms pulse length. Right, representative optical fiber placement. Scale bar = 400 μm (J) Drug-naïve, no interaction: F_(2,20)_ = 0.6640, p = 0.5258. 1 week withdrawal, interaction: F_(2,20)_ = 5.428, p = 0.0131; eYFP vs. Chr2 during on epoch, p = 0.0039. 2-way RM ANOVA followed by Šídák’s multiple comparisons. (K) Cumulative time of nose-to-nose interactions. 2-way RM ANOVA followed by Šídák’s multiple comparisons; interaction: F_(1,10)_ = 7.005, p = 0.0244; eYFP vs Chr2, p = 0.0148). All data are represented as mean ± SEM. #p < 0.1, *p < 0.05, **p < 0.01, ***p < 0.001, ***p < 0.001, ****p < 0.0001.

We first evaluated feeding behaviors, as previously published studies showed that chemogenetic increases of Gi signaling in LS-*Nts* neurons promotes overeating^21,32^. Here, we showed that silencing LS-*Nts* neurons modulated food consumption, but not in the context of opioid withdrawal (**Figure S6A-D**). If LS-*Nts* neurons are indeed activated during naloxone-precipitated withdrawal, they may engage behavioral responding during acute opioid withdrawal. Silencing LS-*Nts* neurons did not influence somatic signs of withdrawal (fecal boli and spontaneous jumping^49–51)^, however, indicating that LS-*Nts* are not necessary for the expression of withdrawal signs (**Figure 7C**). Approximately 24 hours following naloxone, we next tested whether mice experienced hyperalgesia via a hot plate assay and found that TetTox mice licked their hind paw or jumped sooner than controls. TetTox mice also jumped more frequently in the hotplate test at 1d following naloxone treatment, suggesting opioid withdrawal decreased their pain threshold (**Figure 7D**). It was possible that silencing LS-*Nts* elevated gross locomotion, but spontaneous movement in an open field was slightly reduced in TetTox mice—not enhanced (**Figure 7E**, **S6E**). It is also possible that LS-*Nts* neurons control general fear and stress expression. However, TetTox mice spent the same amount of time in the center of an open field as controls (**Figure S6F**) and did not demonstrate an opioid-withdrawal-driven change in elevated plus maze (EPM) performance (**Figure S6G-H**). Together, these data suggested that LS-*Nts* suppression selectively exacerbates pain perception and/or expression during opioid withdrawal.

Opioid withdrawal triggers severe social deficits in humans, including social isolation and aggression, which can perpetuate OUD as individuals are less likely to receive the support necessary to abstain from drug misuse^52–54^. The LS is well-implicated in social behaviors, including social interaction, aggression, mating, and pup-rearing^55^, but it is unknown whether LS-*Nts* neurons play a role in withdrawal-induced social deficits. To test this, TetTox and control mice were placed within a 3-chamber apparatus, in which they were presented with an unfamiliar same-sex mouse in one chamber and an unfamiliar object in the other. At 1 week of opioid withdrawal, TetTox mice spent less time in the mouse-paired chamber than controls (**Figure 7F**), indicating that LS-*Nts* activity influences social preference during early opioid withdrawal. The 3-chamber-assay does not allow naturalistic social interactions, so we next performed a resident intruder task (RI), in which a younger, same-sex albino mouse was placed into the resident mouse’s home cage. TetTox treatment increased the frequency of nose-to-nose interactions in male mice but not females, indicating that silencing LS-*Nts* produces a prosocial state in males (**Figure 7G**). To assess whether optogenetic activation of LS-*Nts* neurons suppresses social interaction before and after opioid withdrawal, we next prepared a cohort of male mice that received bilateral optical fiber implants and injections of AAV5-Ef1a-DIO-Chr2-eYFP (Chr2, n = 6) or AAV5-Ef1a-DIO-eYFP (eYFP control, n = 6) into the LS (**Figure 7H**). These mice underwent a 12 min RI session prior to and 1 week following naloxone-precipitated withdrawal, during which they received interleaved presentations of blue light (**Figure 7I**). In brief, optogenetic activation of LS-*Nts* neurons reduced nose-to-nose interactions with intruder mice during opioid withdrawal (**Figure 7J-K**), which was expected as TetTox-mediated silencing of these neurons promoted social investigation. Further, optically activating LS-*Nts* neurons decreased pain expression (**Figure S6I-K**), demonstrating that LS-*Nts* neurons can bidirectionally regulate social and pain-related behaviors during opioid withdrawal. Finally, we evaluated LS-*Nts* projection targets and found that the retrohippocampal area, lateral hypothalamus, nucleus accumbens, medial and lateral preoptic areas, and the bed nucleus of the stria terminalis were among the top targets of LS-*Nts* neurons (**Figure S7**), indicating that LS-*Nts* projection diversity may underlie this population’s role in adaptive behaviors.

## Discussion

Opioid withdrawal is a powerful motivator for relapse, highlighting the importance for understanding how aversion circuitry in the brain is disrupted by chronic opioid exposure. We chose to study the LS because of its role in mediating both hedonic and aversion responses, which are disrupted during withdrawal. Chronic opioid exposure disrupts synaptic connectivity and structural plasticity within neural circuitry supporting reward and aversion processing^56,57^, molding a system that perpetuates drug craving and relapse vulnerability. Until this point, there was no resource for opioid-driven transcriptional adaptations in the LS, which can provide a guide for which cell types are most pertinent to study in the context of limbic dysfunction in OUD. We utilized our snRNAseq guide to establish LS-*Nts* neurons as a novel neural substrate both perturbed by opioid dependence and selectively disrupted by precipitated withdrawal, and ultimately show that these neurons modulate behavioral responding during protracted opioid withdrawal.

### Developing a functional atlas of the LS

LS neurons receive afferents from hippocampal subregions in a spatially-deterministic manner and may be differentially recruited based on context^9,18,33,58,59^. In fact, the LS may be organized as a lateral inhibition network, where extra-septal glutamatergic input drives different LS cell types to compete with one another^17^. This poses a system where certain LS neurons are preferentially recruited during stress and suppress the activity of other LS cell types to gate behavioral output^7,17,18^. The precise mechanisms through which LS cell types are recruited during opioid withdrawal are unclear. Although *Oprm1* was globally expressed in the septum, its expression across cell clusters was not predictive of overall transcriptional perturbation during chronic morphine treatment. Admittedly, our snRNAseq data do not distinguish among *Oprm1* splice variants or detect protein-level regulation of Mu opioid receptor, which is complex^16^. However, our analyses suggested that the primary driver for LS opioid-induced DEGs may be inputs into the LS. Chronic morphine globally upregulated genes involved in postsynaptic neurotransmission, consistent with what has been reported in other brain areas^38,60^. Furthermore, cell type-specific DEGs were enriched for genes encoding proteins that regulate excitability and synaptic transmission, which may reflect alterations in input-output connectivity. To form predictions about which LS cell types are preferentially engaged during morphine dependence and withdrawal, we performed IEG, DEG and classifier-based analyses across septal clusters. Indeed, opioid dependence perturbed several cell clusters, including LS-*Nts* (Gaba8), LS-*Sst* (Gaba11), LS-*Foxp2* (Gaba7), and glutamatergic neurons in the SFi (Glu2)— which may together form an intra-septal network that facilitates the emergence of opioid-modulated behaviors. To our surprise, however, naloxone-precipitated withdrawal robustly induced gene expression changes in primarily LS-*Nts* neurons (**Figure 3C-E**). It is possible that naloxone-induced LS-*Nts* activation rapidly suppresses the activity of adjacent septal neurons, such as LS-*Esr1*, which showed a transcriptional signature consistent with delayed activation following naloxone treatment. Although we do not directly examine intra-septal connectivity, our snRNAseq data are a comprehensive resource for forming testable predictions of how molecularly-distinct LS cell types communicate with one another.

### The role for LS-*Nts* neurons in behavioral adaptation during opioid withdrawal

Using *in situ*, *in vivo*, and *ex vivo* approaches, we demonstrated that spontaneous withdrawal induces LS-*Nts* activity *in vivo,* likely due to opioid-driven increases in glutamatergic signaling onto LS-*Nts* neurons. Opioid withdrawal is arguably a chronic stress state^1,3,22,61^, and other states of chronic distress, such as pain and chronic social defeat, elevate neuronal activity in the LS, including in LS-*Nts*, LS-*Sst*, and general LS GABAergic neurons^34,62,63^. We originally hypothesized that prolonged LS-*Nts* activation would be maladaptive, but our results indicated the opposite. LS-*Nts* neurons promote analgesia, modulate risk-raking behaviors, and alter how mice approach unfamiliar conspecifics, which could have different consequences for males and females. In other words, LS-*Nts* hyperactivity may drive a series of behavioral responses that are adaptive when the animal is in a weakened, compromised, or uncertain state. Although chronic stress states may be driven by different stimuli (e.g. pain, emotional stress, or drug withdrawal), there is a convergent mechanism that drives LS activity upwards in chronic stress settings. Our snRNAseq data revealed that LS-*Nts* neurons are enriched for genes encoding receptors for neuromodulators, including *Drd2* and *Adra1a*, which have both been extensively studied in the context of drug craving and relapse^64,65^. These observations form rationale for studying how opioid withdrawal disrupts input-regulation onto LS cell types and highlights the potential role for glutamtergic and dopaminergic/noradrenergic signaling as a common driver of LS-*Nts* activity across multiple stress modalities.

LS-*Nts* neurons may encompass more granular cell types not closely studied here. Our snRNAseq and HCR data indicated that *Nts* expression overlaps with *Sst, Foxp2*, *Drd3*, *Met*, and *Crhr2*, which are molecular markers that form layers within the septal complex. The gradient-like cytoarchitecture of the septum poses opportunities for intersectional genetic tools to dissect the precise behavioral properties of each lamina. Given that LS-*Nts* neurons are involved in diverse behaviors, smaller subtypes may comprise the greater *Nts*+ population in the septum and perform distinct behavioral functions. LS-*Nts* neurons also project to diverse brain areas, as shown here and in other studies^7,66,67^, which may be reflected in their molecular diversity. Therefore, future work should evaluate how these neurons modulate downstream targets to gate precise behavioral responding—particularly in the context of opioid-seeking and relapse.

The LS is a unique and heterogeneous limbic structure that performs a diverse role in motivated behaviors, but the precise LS cell types involved in opioid withdrawal were unknown. To address this, we generated the first functional atlas of the septum in opioid dependence and discovered that LS-*Nts* neurons are selectively recruited during opioid withdrawal. Using genetically targeted tools and a combination of *in situ*, *ex vivo*, and *in vivo* approaches, we present here the first demonstration of the multifaceted role for LS-*Nts* neurons in behavioral adaptation during opioid withdrawal. Together, our results emphasize the LS as a key region disrupted by opioid dependence and implicate a new potential therapeutic target in persons with OUD, and our snRNAseq data will provide a comprehensive resource for opioid-perturbed transcriptional alterations in the septum, revealing ample opportunity for future targeted studies on molecularly defined septal neurons.

## Methods

### Animals

Mice were housed in a 12-hour reverse light-dark cycle and provided *ad-lib* access to food and water unless otherwise noted. Both male and female mice were used in all experiments. P56-P63 mice were used in RNA-sequencing and *in situ* experiments, whereas P60 – P180 were used for *in vivo* imaging, *ex vivo* electrophysiology, and behavioral phenotyping. For all experiments, either wildtype C57BL/6J mice (Jackson Laboratories) or *Nts*-Cre mice (Gifted from Zweifel & Palmiter labs, Jackson Laboratory strain #017525) were used. All experiments were conducted in accordance with the National Institutes of Health Guide for the Care and Use of Laboratory Animals and were approved by the Institutional Animal Care and Use Committee and the University of Washington.

### Morphine injections and precipitated withdrawal

To model opioid dependence, we used pharmaceutical-grade morphine procured from UW drug services. Mice were given an escalating dose of morphine over the course of 7 days, with each day successively increased by 10 mg/kg (10 mg/kg on day 1 through 70 mg/kg on day 7). Each mouse received two injections i.p. at each dose ∼10-14 hours apart, for a total of 14 injections. Animals assigned to precipitated withdrawal received a 1 mg/kg dose of naloxone in saline (Sigma-Aldrich, #N7758) i.p. ∼1 hour following their final morphine injection. Control animals only received saline.

### Surgeries

Mice were anesthetized using 5% isoflurane and maintained at 0.8-1.5% isoflurane throughout each surgery. Once induced, they were head mounted in a stereotaxic frame (Kopf Instruments), and ophthalmic ointment was applied to their eyes to protect against dehydration. The incision site was injected s.c. with 1 mg/kg lidocaine, and each mouse received 5 mg/kg s.c. of carprofen, a non-opioid analgesic. We used a Nanoject (Drummond) and pulled glass pipettes (Drummond, #3-000-203-G/X; pipettes pulled using Sutter, P-87) to inject virus at a rate of 1-2 nl/sec into the LS. Coordinates and volumes used for each experiment are detailed in the following sections.

### Single-nucleus RNA-sequencing and library prep

A total of 30 wildtype mice (∼8 weeks) were used for single-nucleus RNA-sequencing. Animals were assigned to one of the following groups: saline (saline only), chronic morphine (escalating morphine), naloxone (escalating morphine plus naloxone), acute morphine (saline 13x, one dose of 10 mg/kg morphine) and naloxone only (saline 13x, one dose of 1 mg/kg naloxone). Each experimental group included 3 male and 3 female mice that were pooled into the same reaction.

Each day for 3 days prior to tissue collection, all animals were habituated to the procedure room for ∼2 hours to control for environmentally-induced immediate early gene activity. Our procedure for tissue collection has been detailed previously^30,31^. On the day of sacrifice, mice were sequentially euthanized with sodium pentobarbital (> 90 mg/kg Somnasol) and transcardially perfused with ice-cold NMDG (96 mM NMDG, 2.5 mM KCl, 1.35 mM NaH_2_PO_4_, 30 mM NaHCO_3_, 20 mM HEPES, 25 mM glucose, 2 mM thiourea, 5 mM Na+ ascorbate, 3 mM Na+pyruvate, 0.6 mM glutathione-ethyl-ester, 2 mM N-acetyl-cysteine, 0.5 mM CaCl_2_, 10 mM MgSO_4_; pH 7.35–7.40, 300-305 mOsm, continuously oxygenated with 95% O2 and 5% CO_2_). This NMDG solution was also used for tissue sectioning and for a collection bath. Both the perfusion solution and recovery bath included an inhibitor cocktail containing 500 nM TTX, 10 μM APV, 10 μM DNQX, 5 μM actinomycin, and 37.7 μM anysomycin, as described previously^30^. 250 μm thick coronal sections containing septum were rapidly harvested using a Leica Vibratome (VT1200). The septal complex was then rapidly microdissected, flash-frozen on powdered dry ice, and stored at -80 °C. We intended to reduce nucleus accumbens contamination as much as possible. To this end, our tissue sections primarily ranged from +1.10 mm through 0.0 mm with respect to Bregma. We also excluded the medial septum and diagonal band level to and ventral to the accumbens. Once tissue from all animals were collected and stored, we proceeded with nuclear isolation, a protocol we adapted from previously-published papers^68,69^. The following steps were all performed on ice. Also note that the PBS used here was Ca^2+^ and Mg^2+^ free. For each experimental group, we pooled all mice into the same tube containing a hypertonic lysis buffer (10mM Tris-HCl, 10 mM NaCl, 3 mM MgCl_2_, 0.1% Igepal, 0.2 U/μl Lucigen NxGen Rnase inhibitor). Nuclei were extracted using a glass dounce homogenizer (DWK, #885300-0002) via 5-10 reps of pestle A and 5 reps of pestle B. Nuclei were then incubated in lysis buffer for 5 min. Next, the nuclear extract was moved into a clean tube, diluted to ∼5-6 ml using a wash buffer (1X PBS, 1% BSA, 0.2 U/μl Rnase inhibitor), and strained into a new 50 ml conical tube using a 40 μm cell strainer (pluriSelect, # 43-50040-51). Strained nuclei were centrifuged for 10 min, 500 x g at 4 °C. The resulting pellet was then retained. To purify our nuclei, we used density gradient centrifugation, where we prepared a two-layer column containing a 25% and 30% iodixanol (OptiPrep, Sigma-Aldrich, #D1556) gradient. Briefly, we suspended the nuclear pellet in a homogenate solution (0.25M Sucrose, 25 mM KCl, 5 mM MgCl_2_, 20 mM Tris-HCl, pH 7.8-8.0, and 0.2 U/μl Rnase inhibitor), and combined the resuspended pellet 1:1 with a 50% iodixanol solution (50% OptiPrep, 25 mM KCl, 5 mM MgCl_2_, 20 mM Tris-HCl, 1% BSA, 0.2 U/μl Rnase inhibitor, pH 7.8-8.0) in a clean 15-ml conical tube. This creates a 25% iodixanol layer. Next, we prepare a 30% iodixanol layer (30% OptiPrep, 25 mM KCl, 5 mM MgCl_2_, 20 mM Tris-HCl, 1% BSA, 0.2 U/μl Rnase inhibitor) and underlaid the 25% iodixanol layer with this 30% layer using a 18G needle. This density gradient column was then centrifuged for 15 min, 3000 x g at 4 °C. The resulting purified and invisible pellet was then resuspended in wash buffer. Nuclei were stained with propidium iodide and assessed for yield and quality using a disposable Neubauer hemocytometer (INCYTO, #DHC-N01) on a fluorescent microscope (Zeiss ApoTome.2). Final nuclear concentration was adjusted to 1,000 nuclei per μl in preparation for single nucleus capture.

Note that the discrepancy in kit version and sequencing platform may have obstructed our ability to compare the newer samples (acute morphine and naloxone only) to the older samples (saline, chronic morphine, withdrawal). Single nuclei were captured, and experimental libraries were prepared using 10X Genomics Chronium Single Cell 3′ V3 Kit (older samples) or 10X Genomics Chromium 3’ NextGEM kit (newer samples). We followed the manufacturer’s protocol and recommendations. Briefly, we captured ∼10,000 barcoded nuclei for each experimental group and reverse-transcribed captured mRNAs to cDNA. cDNA libraries were PCR amplified, fragmented and ligated with sequencing adaptors. cDNA libraries from the older samples were sequenced on the Illumina 4400 HiSeq platform, whereas the newer samples were sequenced on the NovaSeq PE150 platform. Each sample was sequenced to a target output of ∼350,000,000 reads.

### HCR procedure

4 male and 4 female mice were anesthetized with isoflurane and rapidly decapitated. Brains were flash-frozen within powdered dry ice and stored at -80 °C. We collected 20 μm coronal sections directly onto glass slides using a Leica CM3050S, and each slide possessed LS-containing sections from ∼+1.10 through ∼+0.0 Bregma. 1 slide per animal underwent HCR. We used both standard and custom-designed probes provided by Molecular Instruments, and we followed their protocol with some modifications described in our previously-published work^30^. Probes used for this experiment included *Nts*, *Sst*, *Crhr2*, *Drd2*, *Drd3*, *Foxp2*, *Esr1*, *Met*, *Tacr1*, *Pax6*, *Onecut2*, *Samd3*, *Slc32a1*, and *Slc17a6*. In brief, we fixed tissue sections with 4% PFA for 30 min, and performed successive dehydration steps using 50%, 75% and 100% x 2 EtOH. Protease IV (ACDBiosystem) was used to digest tissue for 5 min, and probes were hybridized to tissue at 37 °C. We quenched autofluorescence (Vector labs #SP-8400) and mounted using VectaShield Vibrance antifade mounting media. We acquired images on a Zeiss ApoTome.2 at 10x magnification using Zen (Blue edition). Following each round, tissue was treated with DNase I (Sigma-Aldrich, #4716728001) to deplete the previous round’s probes. To register all images for each section onto the same plane, we took brightfield images, which acted as the reference for each round. To validate the enrichment of *Fos* in LS-*Nts* and LS-*Met* cells following naloxone-induced withdrawal, we performed HCR on tissue from 4 mice treated with saline, 4 mice treated with chronic morphine, and 4 treated with chronic morphine plus naloxone. Each group contained 2 males and 2 females, and every slide contained 1 male and 1 female from the same group (2 slides per group, 4 mice per group). We were interested in the expression of both *Met* and *Nts*, which each ranged throughout the bulk of the septal complex, but we were limited in the number of sections we could test. Therefore, we collected serial sections from ∼+0.85 through ∼+0.0 Bregma, and 2 sections per mouse were retained for analysis. We imaged these sections at 20x magnification to resolve puncta.

### *In vivo* two-photon imaging

4 male and 2 female *Nts*-Cre mice were unilaterally injected in the LS in two positions (rostral: +0.8 AP, - 0.35 ML, -3.0 DV; caudal: +0.2 AP, -0.35 ML, -3.0 DV) with 500 nl of AAVDJ-EF1a-DIO-Gcamp6s (UNC Vector Core, lot #av7830, titer = 2.80 x 10^12^ viral particles per ml). Following injections, a 0.6 mm x 7.3 mm GRIN lens (Inscopix) was slowly lowered and implanted into the intermediate aspect of the LS (+0.5 AP, -0.35 ML, -3.0 DV). We also implanted a metallic ring around the skull for head fixation during imaging. Once animals recovered for 3-4 weeks, we placed them on water restriction to promote motivation for sucrose consumption. For ∼5-7 days, we habituated animals to head-fixation underneath the two-photon microscope, where they intermittently received 10% sucrose boli under the absence of any imaging.

#### Opioid dependence

For Days 0, 2, 4, 6 and 14 (1 week of opioid withdrawal) of the experiment, mice underwent structured imaging sessions. Imaging sessions were designed to first contain a 5 min baseline recording, where the animal remains idle under the 2p, followed by a 30 min session to examine LS-*Nts* evoked activity. For 30 min, mice experienced 150 trials of randomly-delivered 10% sucrose droplets (80% of trials) or 20 psi air puffs directed at the left whisker pad (20% of trials). The ITI varied from 10-15s. Following the first session (Day 0), mice were assigned to either a saline or morphine group (n = 3 for each) and received either saline or morphine injections for 7 days (Days 2, 4, and 6), and were re-imaged again on Day 14 (no opioids). Note that on these days, mice were imaged before they received morphine for the day.

#### Morphine & naloxone-induced changes of activity

Because we were also interested in the acute effects of morphine and naloxone on LS-*Nts* neurons, we designed the imaging session on Day 7 differently than the other days. Mice were first injected with either saline or 70 mg/kg of morphine. ∼30 min following either saline or morphine, each mouse underwent a 5 min baseline recording and a 20 min session, where they received 100 presentations of 10% sucrose and 20 psi air puffs. Upon completion, all mice were injected with 1 mg/kg of naloxone and immediately imaged for an additional 5 min baseline and 20 min task.

All imaging was acquired using an Olympus multiphoton microscope equipped with resonant scanners and GaAsP PMTs as described previously^31,70^. We used a Mai-Tai Deep See laser system (Spectra Physics) at 920 nm wavelength for 2-p excitation. Data were collected at 7 Hz with Olympus FluoView software. Prior to each imaging session, FOVs were manually aligned with averaged projections from the previous day. Once experiments were completed, animals were euthanized using sodium pentobarbital (> 90 mg kg^-1^ Somnasol) and transcardially perfused with (1) 1x PBS and (2) 4% PFA in PBS. Brains were post-fixed for up to ∼2 weeks in 4% PFA to ensure lens stability in the brain. Brains were next moved to 30% sucrose in 1x PBS for 2 days, and cryo-sectioned at 40 μm on a Leica CM3050S. To enhance and stabilize Gcamp6s signal, sections were blocked, permeabilized and then stained with 1:500 chicken anti-GFP antibody (Aves Labs, #GFP-1020) overnight at 4 °C and then with 1:1000 donkey anti-chicken 488 (Jackson ImmunoResearch, #703-545-155) for 2 hours at RT. Sections were then imaged on a confocal microscope (Olympus FV 3300).

### *Ex vivo* electrophysiology

All electrophysiology experime*nts* were performed on brain slices collected at approximately the same time of day. Mice were euthanized with an i.p. injection of sodium pentobarbital (> 90 mg/kg Somnasol). Mice were decapitated and the brain was extracted. After extraction, the brain was immersed in ice-cold NMDG ACSF (92 mM NMDG, 2.5 mM KCl, 1.25 mM NaH_2_PO, 30 mM NaHCO_3_, 20 mM HEPES, 25 mM glucose, 2 mM thiourea, 5 mM Na-ascorbate, 3 mM Na-pyruvate, 0.5 mM CaCl_2_·4H_2_O, 10 mM MgSO_4_·7H_2_O, and 12 mM N-Acetyl-L-cysteine; pH adjusted to 7.3-7.4) for 2 min. Afterwards coronal slices (300 µm) were sectioned using a vibratome (VT1200s, Leica, Germany) and then incubated in NMDG ACSF at 34 °C for approximately 14 min. Slices were then transferred into a holding solution of HEPES ACSF (92 mM NaCl, 2.5 mM KCl, 1.25 mM NaH_2_PO_4_, 30 mM NaHCO_3_, 20 mM HEPES, 25 mM glucose, 2 mM thiourea, 5 mM Na-ascorbate, 3 mM Na-pyruvate, 2 mM CaCl_2_·4H_2_O, 2 mM MgSO_4_·7H_2_O and 12 mM N-Acetyl-l-cysteine, bubbled at room temperature with 95% O2/ 5% CO2) for at least 45 mins until recordings were performed.

Whole-cell recordings were performed using a Multiclamp 700B (Molecular Devices, Sunnyvale, CA) using pipettes with a resistance of 4-6 Mohm. Pipettes were filled with an internal solution containing 100 mM cesium gluconate, 0.6 mM EGTA, 10 mM HEPES, 5 mM NaCl, 20 mM TEA, 4 mM Mg-ATP, and 0.3 mM Na-GTP with the pH adjusted to 7.2 with CsOH and the osmolarity adjusted to around 289 mmol/kg with sucrose. During recordings, slices were perfused with a recording ACSF solution (120 mM NaCl, 3.5 mM KCl, 1.25 mM NaH_2_PO_4_, 26 mM NaHCO_3_, 1.3 mM MgCl_2_, 2.4 mM CaCl_2_ and 11 mM D-(+)-glucose, and was continuously bubbled with 95% O2/5% CO_2_). Infrared differential interference contrast–enhanced visual guidance was used to select neurons that were 3–4 cell layers below the surface of the slices. LS-*Nts* neurons were identified by the presence of td-Tomato using a fluorescence microscope (Scientifica SliceScope Pro 1000; LED: SPECTRA X light engine (Lumencor)). The recording solution was delivered to slices via superfusion driven by peristaltic pump (flow rate of 4-5 ml/min) and was held at room temperature. The neurons were voltage clamped at −70 mV or +10 mV, and the pipette series resistance was monitored throughout the experime*nts* by hyperpolarizing steps of -10 mV with each sweep. If the series resistance changed by >20 % during the recording, the data were discarded. Whole-cell currents were filtered at 1 kHz and digitized and stored at 10 KHz (Clampex 10; MDS Analytical Technologies). All experiments were completed within 4 hours after slices were made to maximize cell viability and consistency.

mEPSCs and mIPSCs were recorded in the presence of TTX (1 μM), d-AP5 (50 μM) in the recording ACSF solution. For one example recording, gabazine (1 μM) was added while holding at +10 mV to confirm observed currents were mediated by GABAA receptors. Data was analyzed with Easy Electrophysiology. For mEPSCs, a detection threshold of >10 pA and rise time <3 ms was used; for mIPSCs, a detection threshold of >10 pA and rise time <10 ms was used. Results were visually verified.

To compare the mEPSC and mIPSC frequency and amplitudes across conditions, linear, mixed-effects regressions (LMERs) were performed using the “lmer” function in R. Interevent interval (IEI) data was log-transformed while amplitude data was inverse-transformed to best fit a normal distribution. For statistical analysis of both IEI data and amplitude data, the linear, mixed effects regression model included group (Morphine vs Saline) as a fixed effect and mouse and cell as nested random effects. Because LMERs do not require an equal number of observations from each cell, all events from a given recording were used, with recordings typically containing 100-1000 events. Effect sizes were calculated as Cohen’s d using the “lme.dscore” function in the “EMATools” package in R.

### TetTox & Optogenetics behavioral profiling

For TetTox experiments, 14 *Nts*-Cre mice (6 males, 8 females) and 12 Nts-Cre mice (6 males, 6 females) were bilaterally injected with 400-500 nl of AAVDJ-EF1a-DIO-TetTox:GFP (Gifted from Palmiter lab, titer = 4.2 x 10^12^ viral particles per ml) <u>or</u> AAV5-EF1a-DIO-eYFP (UNC Vector Core, lot #av4802B, titer = 4.40 x 10^12^ viral particles per ml) in the LS (0.6 AP, ± 0.40 ML, -2.80 DV), respectively. The virus was allowed to incubate for at least 8 weeks before any behavioral testing. Following this recovery period, baseline testing occurred over 1-2 weeks. For optogenetics, 12 male *Nts*-Cre mice were bilaterally injected with 400 nl of AAV5-Ef1a-DIO-hChr2-eYFP (UNC Vector Core, AV43132) or AAV5-Ef1a- DIO-eYFP (UNC Vector Core, AV4310L) into the LS (0.38 AP, ± 0.45 ML, -2.90 DV), and received bilateral implantation of optical fibers (RWD Life Science, Inc; 907-03007-00) over the same site (10°, 0.38 AP, ± 1.01 ML, -2.86 DV). Mice were allowed to recover for ∼4 weeks prior to baseline testing. All mice underwent morphine escalation as described in other experiments. On completion of the drug course, behavioral testing resumed for up to 4-5 weeks. Prior to each day’s testing, all mice were acclimated to the room for 30 min – 1 hour. Recordings were captured under an infrared light source and tracked via Noldus Ethovision XT11, unless otherwise noted.

#### Open field

Mice were allowed to explore a standard open field for 10 minutes and were recorded from overhead. The first 5 minutes of data were retained for analysis. For optogenetic experiments, laser light was *Elevated plus maze*. Mice were allowed to explore a standard EPM setup (13.5 in height, 25 in length of each arm, 7 in high wall on closed arms) for 10 min. Annotated zones included open and closed arms, as well as the center of the EPM. The first 5 minutes of data were retained for analysis.

#### Sucrose preference

We used Med Associates chambers and voltage lickometers to assess each animal’s preference for tap water or 2% sucrose solution. Mice were habituated to Med Associates chambers for 2-3 sessions, 30 min – 1 hr each, prior to the test session, and test sessions were 1 hr long.

#### Social preference

To test for differences in social proclivities, we used a 3-chamber setup in which subjects were presented with an unfamiliar mouse on one side and an unfamiliar object on the other. Mice were first habituated to the 3-chamber apparatus with nothing in either side, and on the following day, they were exposed to a same-sex, weight-matched mouse and a 3D printed mouse for 10 min.

#### Hot plate

Mice were exposed to a hot plate (BioSeb) for 30 s at 55 °C to evaluate pain-coping behaviors, including hind-paw licks and jumps. Recordings of all animals were coded and blindly scored, and BORIS^71^ was used for behavioral annotation and quantification.

#### Withdrawal signs

To assess differences in withdrawal signs, mice were injected with 1 mg kg^-1^ of naloxone 1 hour following their final dose of morphine. They were immediately placed into an empty cage and recorded from the side for 20 min. Recordings of all animals were coded and blindly scored for withdrawal signs, and BORIS^71^ was used for behavioral annotation and quantification. Withdrawal signs included fecal boli, jumping, and bouts of lethargy.

#### Resident intruder assay

We performed a resident intruder assay ∼2 months of opioid withdrawal. A young (∼12 week) same-sex albino mouse (C57BL/6J) was placed into the home cage of each resident mouse for 6 min, and social interactions were recorded from the side. Videos were scored using BORIS, and we measured the following behaviors of the resident mouse: nose-to-nose sniffing, anogenital sniffing, fleeing, digging, and huddling. Notably, there were no attack bouts.

### SHIELD Tissue clearing

To label LS-*Nts* projections, 4 *Nts*-Cre mice were unilaterally injected with 200 nl of AAV8-EF1a-DIO-eYFP-NRN (Stanford Virus Core, cat# GVVC-AAV-164, 3.08 x 10^13^ viral particles per ml, 1:1 dilution with saline) in the LS (+0.60 AP, 0.40 ML, -2.80 DV). Three weeks following viral injection, mice were perfused with 1x PBS and 4% PFA-PBS, after which the brain was harvested and post-fixed with PFA for 24 hours. After post-fixation, the brains were processed following the SHIELD tissue clearing protocol (LifeCanvas). Briefly, the brain was incubated in SHIELD OFF solution [2.5 mL DI water, 2.5 mL SHIELD-Buffer Solution, 5 mL SHIELD-Epoxy Solution] at 4°C for 4 days. Then transferred to SHIELD ON buffer and incubated at 37°C for 24 hrs. Following, SHIELD ON buffer incubation, the brain was pre-incubated in Delipidization buffer for overnight at room temperature. The tissue was then placed into SmartClear (LifeCanvas) to conduct active tissue clearing for 24 hours. To match the refraction index, the tissue was incubated in 50% of EASI-INDEX solution (RI = 1.52, LifeCanvas) at 37°C for 24 hrs and then 100% of EASI-INDEX solution at 37°C for 24 hrs. The tissue was then embedded in EASI-INDEX agar and mounted onto SmartSpin light sheet microscope (LifeCanvas). Tissue was imaged from the horizontal view using a 3.6x objective with 4 μm zstep. Images were registered to a standard brain atlas (Chon et al., 2019) using ClearMap pipeline (Renier et al., 2016) on a cluster computer. eYFP expressing puncta spots were segmented through the same pipeline using a classifier created using Ilastik (Berg et al., 2019). The total number of eYFP positive puncta spots per brain region in the injection ipsilateral hemisphere was quantified.

## Quantification & statistical methods

### General statistics

Statistical methods and results were reported in each figure. Data generated from our TetTox experiments were analyzed using Prism v9.4.1. Generally, we performed repeated measures two-way ANOVA followed by Sidak’s multiple comparisons correction. Some analyses leveraged linear regression to compare y-intercepts and slopes among different groups and timepoints, where noted. Sequencing and calcium imaging data were analyzed using both published pipelines and custom Python and R scripts.

### Single-cell RNA-sequencing clustering and gene expression analysis

Once sequenced, reads were aligned to a pre-mRNA reference genome using the 10X Genomics’ Cell Ranger v3 pipeline, which provided digital expression matrices that were used for downstream analysis. Clustering, sample integration, cell type identification, and differential gene expression identification were all carried out using the Seurat V3 R package^72^.

#### Filtering and quality control

We have previously published our general filtering parameters^30,31^, but in brief, genes expressed in fewer than 3 cells were removed from the dataset. We next retained cells that possessed between 500 and 25,000 UMIs and fewer than 1% mitochondrial read enrichment. Although we included the cutoffs normally used for scRNA-seq, snRNAseq data are mostly depleted of mitochondrial reads. Doublets were then computationally predicted and removed by the DoubletDecon R package^23^ using the default parameters.

#### Sample integration and clustering

Once putative doublets were omitted, we utilized Seurat’s integration algorithm based on canonical correlation analysis^24^ and mutual nearest neighbor analysis^25^. Gene transcript counts were scaled for total sequencing depth (factor of 10,000) and then natural-log transformed. Integration anchors were identified for each dataset and transformed by a correction vector to produce a “corrected” expression matrix, on which we performed principal component analysis (PCA). We used the first 30 PCs to identify cell clusters (using Louvain, resolution 0.80) and to plot their proximity in space using Uniform Manifold Approximation and Projection (UMAP). We evaluated the expression of canonical neuronal markers (e.g. *Stmn2* and *Thy1*) to identify neuronal populations, and proceeded to subset these neurons out of the data and performed re-clustering to reveal intra-neuronal heterogeneity. In re-clustering neurons, we normalized and scaled expression data using scTransform^73^, which constructs a generalized linear model (GLM) to dissociate sequencing depth from gene expression without reducing cell type heterogeneity. We integrated datasets and clustered individual neurons based on these transformed data as described before, but used the original, linearly-scaled and normalized RNA values in downstream expression analyses.

To determine the optimal clustering resolution for our data (i.e. number of bioinformatically-determined neuron types), we utilized the Clustree R package^74^. This approach allowed us to assess the stability of clusters over a series of clustering granularities. We determined 14 clusters (2 glutamatergic and 12 GABAergic at a resolution of 0.20) to be optimal because it produced stable clusters with neurons yielding from each experimental group. Although it appears that perhaps a lower resolution would produce even more stable clusters (resolution = 0.10), we selected the higher value because we believe that Gaba12 neurons are bona-fide septal cells, distinct from Gaba7. They’re however rare, and so do not separate as readily as other cells might. Next, to identify potential molecular markers for each cell cluster, we computed the average expression of a given gene in one cluster, and then compared it to the expression of that gene in all other clusters combined. P-values were computed using Wilcoxon rank sum test and corrected for the total number of genes tested.

#### DEG analysis

To understand what transcriptional changes were induced by chronic morphine treatment, we first collapsed cell types into single pools separated by condition (Sal, Mor, and Nal). We did this first because presumably the most robustly altered DEGs would be enriched in pseudobulk. We performed a series of pairwise comparisons across all genes between two experimental groups (Sal vs Mor; Mor vs Nal) and retained genes with an average logFC of at least 0.1 and p-value < 0.05 (Wilcoxon rank sum test following by Bonferroni post-hoc correction). We then took this list of pseudobulk DEGs and split them as either upregulated (logFC > 0.1) or downregulated (logFC < -0.1) and fed these lists through EnrichR for GO analysis^75–77^. We referred to the top 10 GO terms to derive a list of glutamate receptors, GABA receptors, potassium channels, and calcium channels, and we subsequently calculated the average logFC between Mor and Sal, such that positive logFC values indicated enrichment in Mor (Wilcox rank sum test followed by Benjamini-Hochberg false discovery rate, adjusted for the number of cell clusters tested; n = 14).

We also wanted to understand which morphine-induced DEGs may change in a cell type-specific manner. To do this, we identified Mor-induced DEGs for each cell cluster because we wanted to capture DEGs that may be uniquely altered on a cell cluster basis. We then concatenated the DEGs identified from every cell cluster and took the fold-change difference between these two datasets, resulting in a Δ expression matrix for saline versus morphine. Dimensionality of this data frame was reduced via PCA, and the feature ranking (genes) from PC1 was retained and plotted as a heatmap. We identified enriched ontology terms using EnrichR. Note that we removed Gaba12 from this analysis because it was transcriptionally distinct from the other cell types, thereby skewing the variability of DEG expression.

To determine a DEG enrichment score for Sal vs Mor, we randomly sampled at most 100 cells from each neuron cluster and summed both up- and down-regulated DEGs into one value. This is because there is an innate correlation between the number of cells per cluster and the number of DEGs that will be identified; limiting the number of tested cells for each cluster limits this bias. These values were normalized across cell types in each comparison, such that the lowest DEG total was 0 and the highest was 1. We approached the Mor vs Nal comparison differently because gene expression differences were much more subtle these two groups. To ensure our DEG metric for Mor vs Nal was stable, we randomly sampled 100 cells from each cell type and each group to compute the total number of DEGs and repeated this 100 times. We took the normalized average number of DEGs as the score for Mor vs Nal.

#### IEG analysis

We used a predetermined list of IEGs implicated in the literature^27^. From this list, we first subset genes that were present in our LS data and calculated the average expression difference between Mor and Nal of each gene in each cluster and subsequently tested whether the difference in expression between Mor and Nal was significant (Wilcox rank sum test followed by Benjamini-Hochberg false discovery rate, adjusted for the number of cell clusters tested; n = 14). In Figure 3, we plot only the genes that were significantly altered by precipitated withdrawal in any cell cluster. To determine a singular metric for neuronal activation post-naloxone, we first averaged the expression of the panel of significantly altered IEGs within each cell type (single IEG average for every cell), then subsequently tested whether this average was statistically different between Mor and Nal conditions (Wilcox rank sum test followed by Bonferroni post-hoc correction). We performed the same analysis with *Fos*, *Fosb*, *Egr1*, *Jun*, *Junb, Jund, Arc,* and *Nr4a1*.

#### Classifier analysis

We performed pairwise comparisons between Sal and Mor and then Mor and Nal to compute AUC, a measure of classifier accuracy, using the R package Augur^42^ with the default settings (Random Forrest, 20 randomly-sampled cells across 150 restarts). This analysis was rooted in the assumption that the classifier will be more accurate if cells between each condition are more transcriptionally distinct. Lower AUC scores indicate that cells between two conditions are more similar. AUC scores across 150 iterations were plotted for Sal vs Mor and Mor and Nal and statistically tested for differences against a shuffled control (Sal vs Mor, but group labels for each cell were randomized).

### HCR analysis

All images acquired for HCR were registered and cropped using ACDBiosystem’s HiPlex image registration software, with brightfield images used as a reference. We used Indica Labs HALO v3.2 to perform multi-channel *in situ* spot quantification. Parameters for detecting puncta and intensity of each gene were manually adjusted for each tissue section of each animal. Cells were considered positive for a particular gene if they possessed 5 or more copies. Cell type abundance and spatial distributions were calculated using custom R scripts.

#### Correlation between snRNAseq and HCR

To evaluate the congruency between snRNAseq and HCR datasets, we first clustered the *in situ* data. We performed supervised clustering by feeding the HCR dataset into Seurat as a digital expression matrix. This HCR-based expression matrix contained single cells detected across all 8 animals and the spot counts for every gene; we also included signal intensity measurements as metadata. Once loaded, the HCR digital expression matrix was scaled but not normalized. During scaling, we regressed out gene intensity measurements on the basis that signal intensity may indicate staining quality. Next, cells were clustered using the same pipeline as described above. We manually identified which HCR cluster corresponds to each snRNAseq cluster based on relative expression of predicted cell type markers. We computed the percent of cells expressing every candidate gene in each cluster and correlated gene enrichment between HCR and scRNAseq datasets via Pearson correlation.

#### Calculating naloxone-induced IEG expression changes

To measure *Fos* induction in LS- *Nts* and LS-*Met* neurons, we collapsed cells from all animals in each experimental group. The experimenter adjusting HALO detection parameters for every gene was blinded to the experimental group of each tissue section. Any cell containing the top 15^th^ percentile of spot counts for *Nts* and *Met* were determined to be *Nts*+ or *Met*+, respectively. However, puncta for IEGs did not undergo thresholding and were instead treated as distributions. Once spot counts were determined for all sections and animals, 2-3 sections per animal were retained for analysis, and all cells from all animals were collapsed into the same dataset. Determining whether Nal induces IEG expression required us to perform a linear mixed effects model to account for variance associated with animal and sex (fos.Copies ∼ group + (1 | sex) + (1 | slide) + (1 | animal)). We also collapsed all cells into Sal, Mor and Nal groups and compared relative distributions of each gene within each cell type using the Kolmogorov-Smirnov test (CDF included 10 bins). We chose not to run statistical testing on the median expression of each gene (of collapsed cell data) because the n was high enough to induce significance in any comparison.

### Two-photon analysis

Raw calcium imaging files were exported as a series of .TIFF images and loaded into Suite2p^78^ for motion correction, ROI detection and signal extraction. Motion correction was performed using nonrigid registration with a 300-frame averaged reference, and an integrated CellPose^79^ plugin was used to automate ROI detection. We manually annotated cells that were not detected. Due to high correlation between raw fluorescence traces and output neuropil signals, we elected to not perform neuropil correction and instead retained only the raw fluorescence trace for downstream analysis. We were concerned that because LS-*Nts* are relatively sparse, the neuropil signal may be contaminated by the cell itself.

#### Spontaneous activity analysis

Assessing withdrawal-induced alterations in spontaneous activity required us to detect discrete calcium events in our data. To achieve this, we wrote a custom Python script to perform the following transformations to each calcium trace: (1) rolling mean over 20 frames, (2) baseline correction using a polynomial fit, (3) Z scoring across the trace, (4) Buttersworth lowpass filter in both forward and reverse directions to correct for time lag, and (5) peak detection via SciPy Signal package. Due to preprocessing and filtering, we did not threshold peaks by amplitude in the final step.

#### Evoked activity analysis

Within each calcium trace, we identified epochs when (1) sucrose or (2) air puffs were delivered and identified peri-stimulus time histograms (PSTHs) for every event, similar to how we have reported before^70^. When calculating ΔF/F, we used the 2.5 s period between the trial start and stimulus delivery as a the F_0_ (baseline). For reported Z scores, we performed Z scoring across the mean PSTH for every cell. To determine whether a cell is responsive to each stimulus, we used a Wilcoxon signed rank test on its baseline activity (F_0_) versus its evoked activity on every trial. Multiple comparisons for each cell were adjusted using Bonferroni post-hoc correction. We then counted the number of cells deemed responsive (p < 0.05) and performed Fisher’s Exact Test to determine whether morphine dependence alters LS- *Nts* responsivity to sucrose and air puffs. Finally, to compute AUC, we used a linear trapezoidal method over 5 s following stimulus presentation.

## Acknowledgements

We thank the lab of Richard D. Palmiter for the TetTox virus and the lab of Larry S. Zweifel for the Nts-Cre line. This work was supported by the University of Washington Addictions, Drug & Alcohol Institute (UW ADAI) (R.C.S.) and NIH grants R37 DA032750 (G.D.S.), R01 DA038168 (G.D.S.), and F31 MH117931 (R.C.S.). This work was also supported by the University of Washington Center of Excellence in Neurobiology of Addiction, Pain and Emotion (NAPE Center, P30 DA048736).

## Author Contributions

Conceptualization, R.C.S and G.D.S.; Methodology, R.C.S. and G.D.S.; Validation, R.C.S. and B.A.B.; Formal analysis, R.C.S., P.S., W.T.F., K.I., and K.H.; Investigation, R.C.S., W.T.F., K.I., M.D.H., M.M.M., B.A.B. and A.D.S.; Resources, G.D.S. and S.A.G.; Data curation, R.C.S. and P.S.; Writing – original draft, R.C.S. and G.D.S.; Writing – review & editing, R.C.S. and G.D.S. with input from all authors; Visualization, R.C.S., P.S., W.T.F., and K.I.; Supervision, G.D.S.; Funding acquisition, G.D.S. and R.C.S.

## Resource Availability

### Lead Contact

Further information and requests for resources and reagents should be directed to and will be fulfilled by Garret Stuber.

### Data and Code Availability

The NCBI Gene Expression Omnibus accession number for the scRNAseq data reported in this paper will be available upon publication. All the code used to analyze scRNAseq, HCR, and two-photon imaging data are available at a Github repository affiliated with Stuber Laboratory group and this manuscript title (http://www.github.com/stuberlab/).

## Supplemental Figures

**Figure S1.**
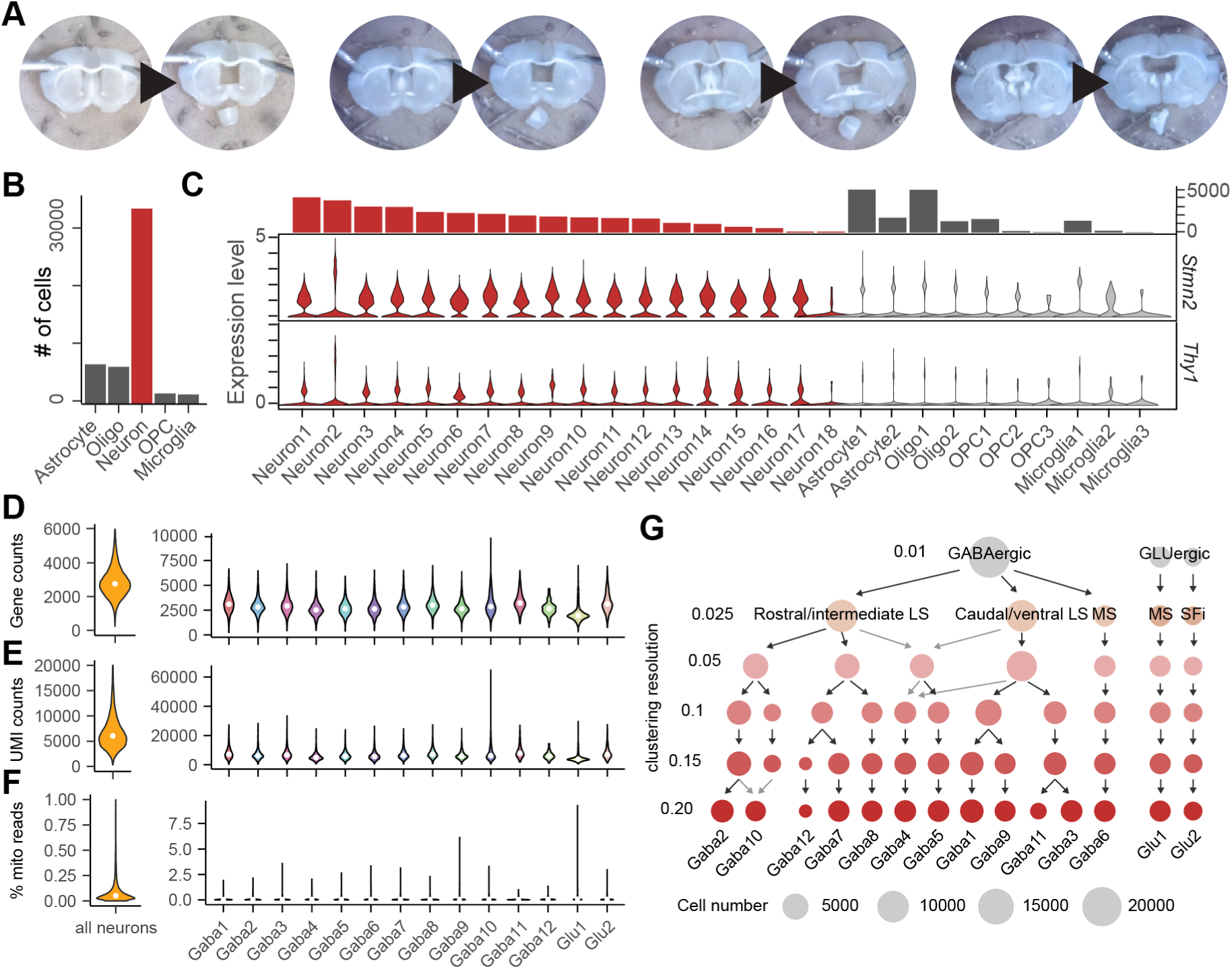
snRNAseq quality metrics and clustering analysis. (A) Sample tissue dissections of the septum. The LS was the primary target and much of the medial septum and diagonal band were excluded. (B) Number of nuclei collected per cell class. (C) Violin plots of *Stmn2* and *Thy1* expression among all cell clusters. Putative neuron clusters are labeled in red. (D) Distribution of features (gene counts) for all neurons (left) and for each neuron cluster (right). (E) Distribution of UMIs (RNAs) for all neurons (left) and for each neuron cluster (right). (F) Distribution of % mitochondrial reads for all neurons (left) and for each neuron cluster (right). (G) Clustree diagram of neuronal clustering across multiple clustering granularities. Disc size indicates the approximate number of cells within each cluster, and color indicates resolution.

**Figure S2.**
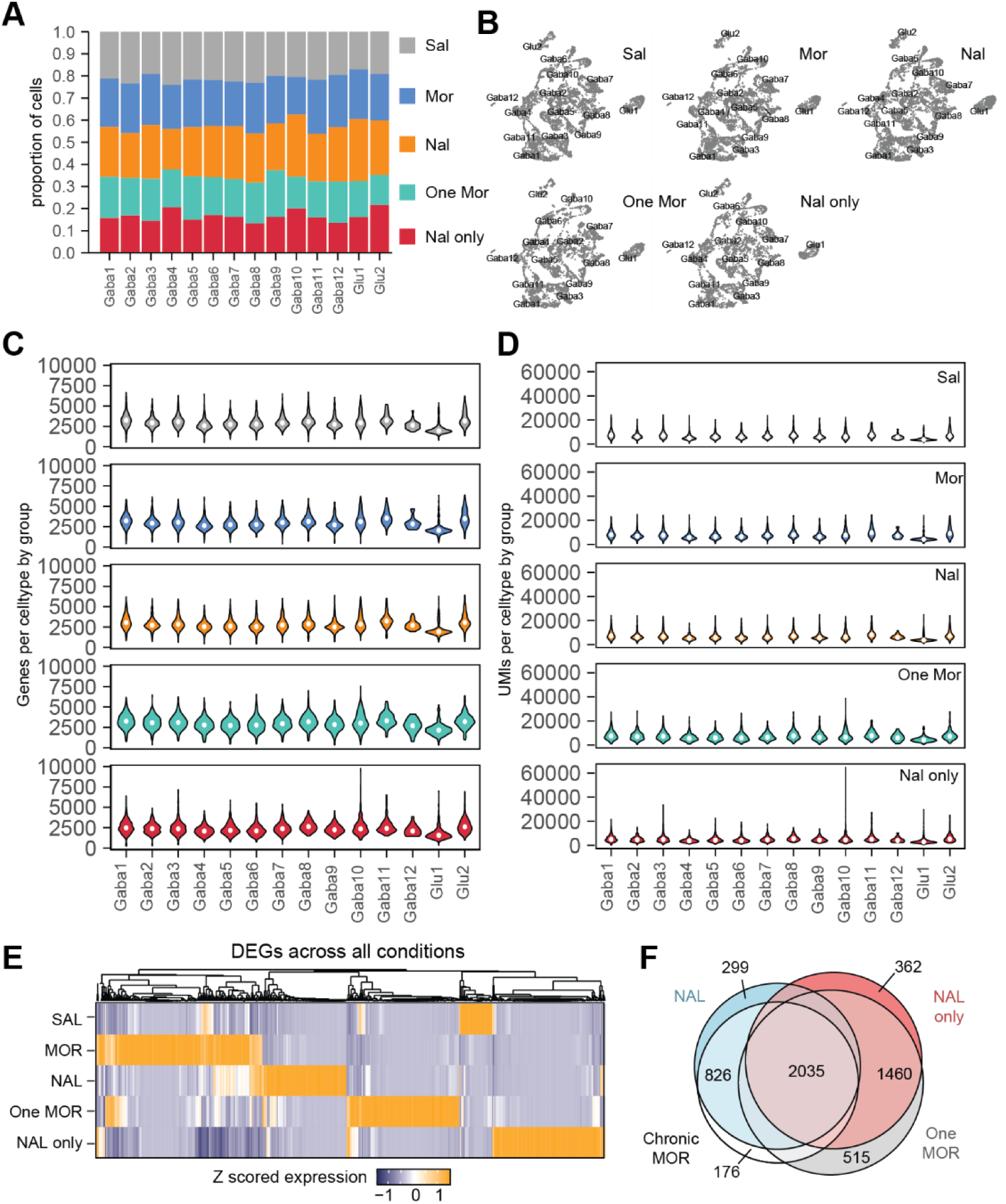
Representation of each experimental group in snRNAseq data. (A) Proportion of cells in each experimental group across neuron clusters. (B) Plots indicating the location of cells assigned to each group in UMAP space. (C) Distribution of features (genes) detected across neuron clusters and experimental groups. (D) Distribution of UMIs (RNAs) detected across neuron clusters and experimental groups. (E) Heatmap of DEG expression (Z scored) for each experimental group identified via pseudobulk. Each group was compared to Sal to identify DEGs. (F) Euler diagram of the overlap of DEGs for each experimental group. Note the extraordinarily high number of DEGs induced by short term treatments.

**Figure S3.**
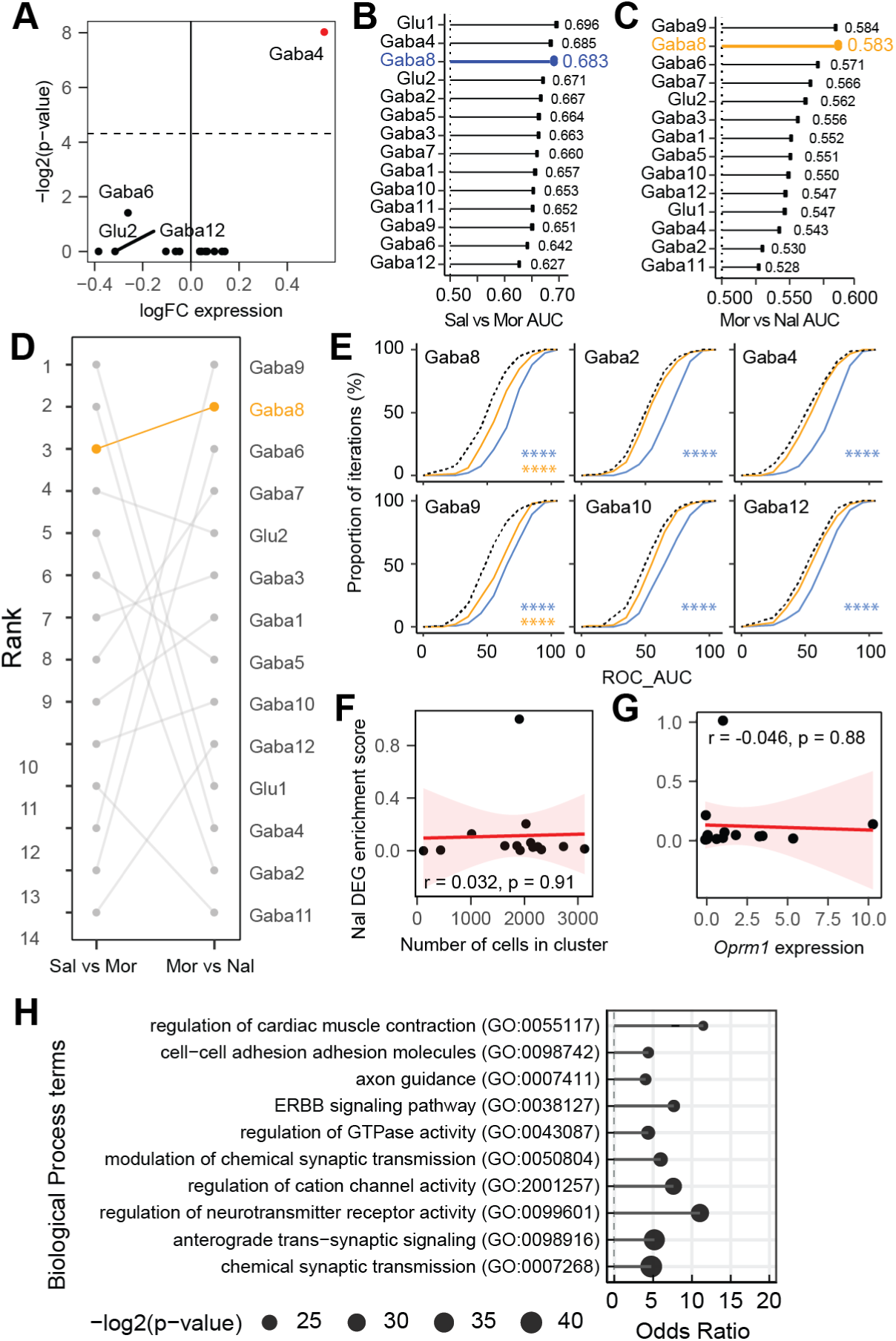
Selecting the most transcriptionally perturbed cell type in the septum. (A) Average logFC expression difference between Mor and Nal of “classic” IEGs for each cell cluster. Wilcoxon rank sum test followed by Bonferroni’s post hoc correction. (B-C) Lollipop plots illustrating computed AUC scores for each cell type for Sal vs Mor (B) and Mor vs Nal (C). A higher AUC score indicates a higher degree of transcriptional change in each condition comparison. (D) Bump plot of AUC score rankings for each cell type across each comparison: Sal vs Mor and Mor vs Nal. (E) Distribution of AUC scores over 150 runs for Gaba8, Gaba9, Gaba2, Gaba10, Gaba4 and Gaba12 in the Sal vs Mor comparisons (blue) and Mor vs Nal comparisons (orange). Shift in distribution was tested using the Kolmogirov-Smirnov test, corrected by Bonferroni’s post hoc. (H) Correlation between the number of cells in each cluster (x-axis) and Nal DEG enrichment score (y-axis). Pearson correlation. (I) Correlation between *Oprm1* expression (x-axis) and Nal DEG enrichment score (y-axis). Pearson correlation. (I) Top 10 Biological Process terms enriched in the genes identified to differentiate *Nts*+ neurons from *Nts*-neurons.

**Figure S4.**
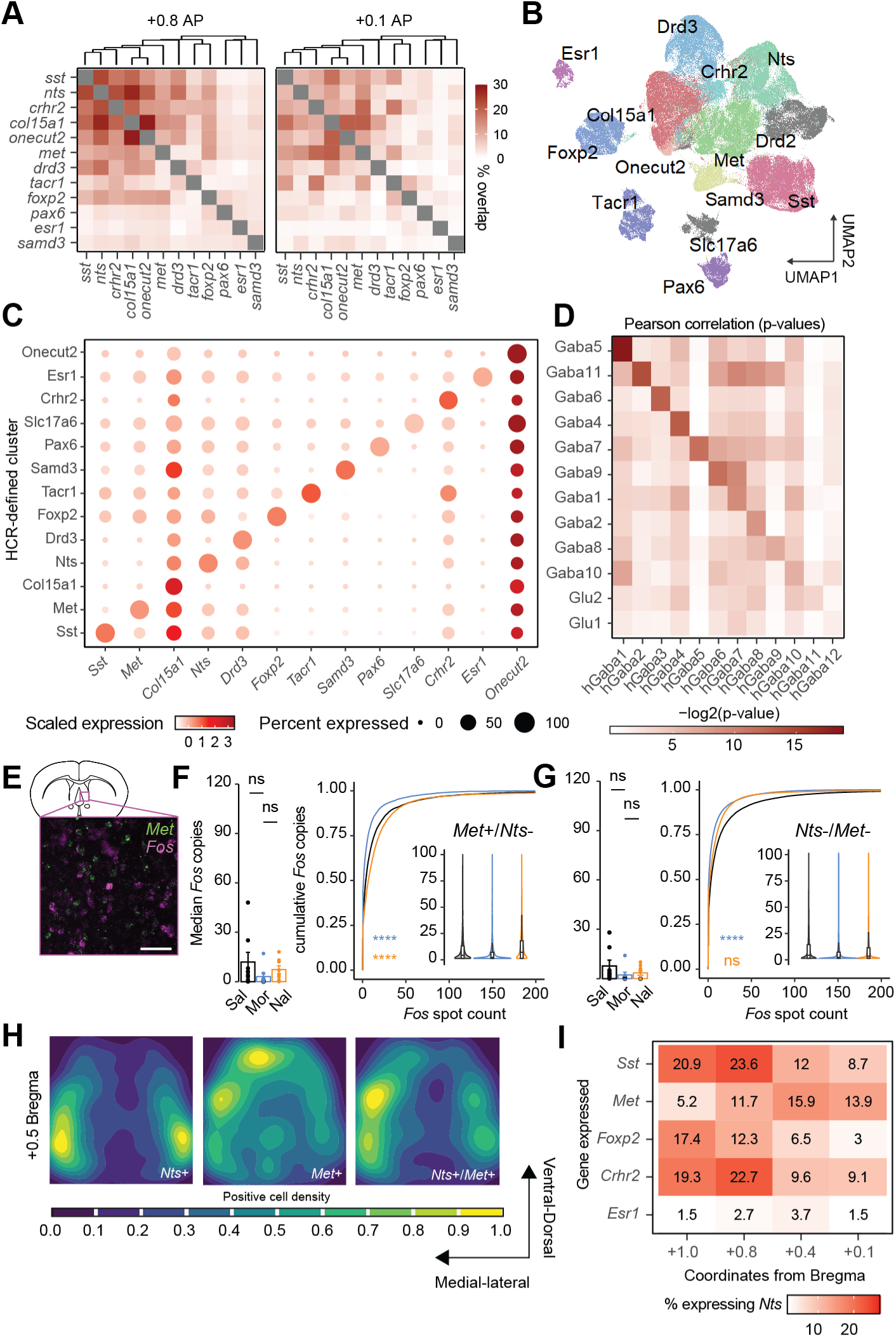
Related to Figure 4, HCR clustering and IEG enrichment. (A) Heatmaps indicating the percent overlap between cells expressing each gene in anterior septal sections (left) and posterior septal sections (right). (B) UMAP plot of cells clustered based on HCR data. Clusters were manually assigned names according to their expression profiles. (C) Disc plot of the expression of each HCR gene in each cluster. (D) Heatmap of p-values corresponding to correlation values in Figure 4E, Right. Pearson correlation. (E) Sample *Met* and *Fos* overlap. Scale bar = 100 μm. (F) Cells positive for *Met* only. Sal vs Mor: median copies per section (p = 0.706); median copy difference (p = 4.62 x 10^-5^). Mor vs Nal: median copies per section (p = 0.127); median copy difference (p = 9.44 x 10^-7^). (G) Cells negative for *Nts* and *Met*. Sal vs Mor: median copies per section (p = 1); median copy difference (p = 2.48 x 10^-10^). Mor vs Nal: median copies per section (p = 1); median copy difference (p = 0.2). (H) Density plot indicating the overlap of *Nts*+ and *Met*+ cells at ∼+0.50 Bregma. Statistics: #p < 0.1, *p < 0.05, **p < 0.01, ***p < 0.001, ***p < 0.001, ****p < 0.0001. (I) Heatmap illustrating the percent of cells that express coexpress *Nts* and a given gene across the AP axis of the LS.

**Figure S5.**
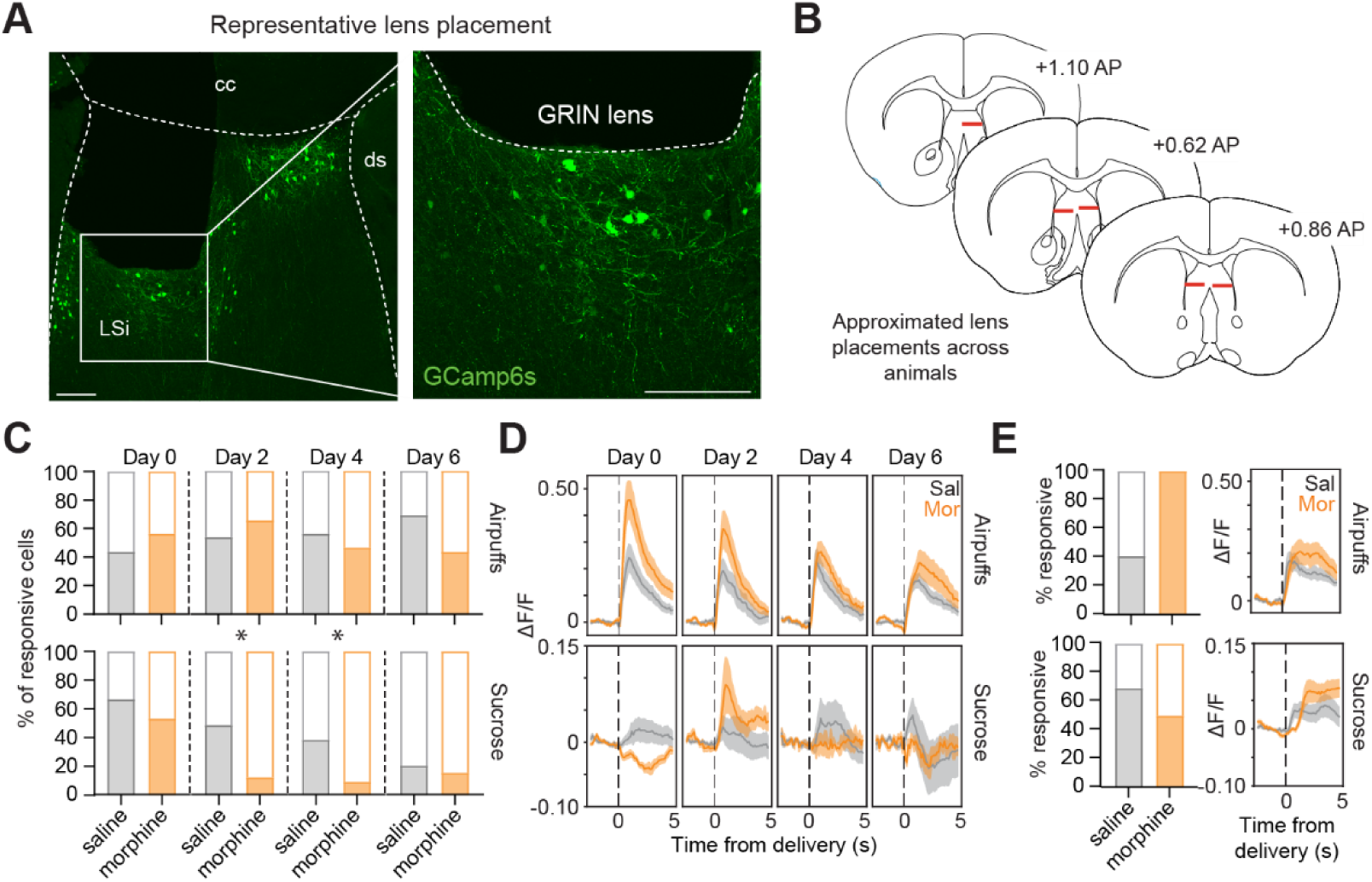
Emergence of opioid dependence in septum. (A) Representative image of GRIN lens placement and GCamp6s expression in the septum. Scale bar = 200 μm. (B) Qualitative mapping of LS GRIN lens placements across 5 animals. One animal was omitted due to incorrect placement. Sal group: n = 3; Mor group: n = 2. (C) Proportion of neurons responsive to air puffs (top) and sucrose presentations (bottom) across Day 0 through Day 6. Significance was computed using Chi Square Test. (D) Mean activation of neurons responsive to air puffs (top) and sucrose presentations (bottom) across Day 0 through Day 6. Significance was computed on the AUC following stimulus presentation using a 2-way repeated measures ANOVA followed by Šídák’s multiple comparisons. (E) Left, proportion of cells responsive to air puffs (top, Sal: n = 39; Mor, n = 32) and sucrose (bottom, Sal: n = 39; Mor, n = 23) on day 14. Right, mean activation by air puffs (top) and sucrose (bottom). No comparisons were statistically significant. All data are represented as mean ± SEM. Statistics: #p < 0.1, *p < 0.05, **p < 0.01, ***p < 0.001, ***p < 0.001, ****p < 0.0001.

**Figure S6.**
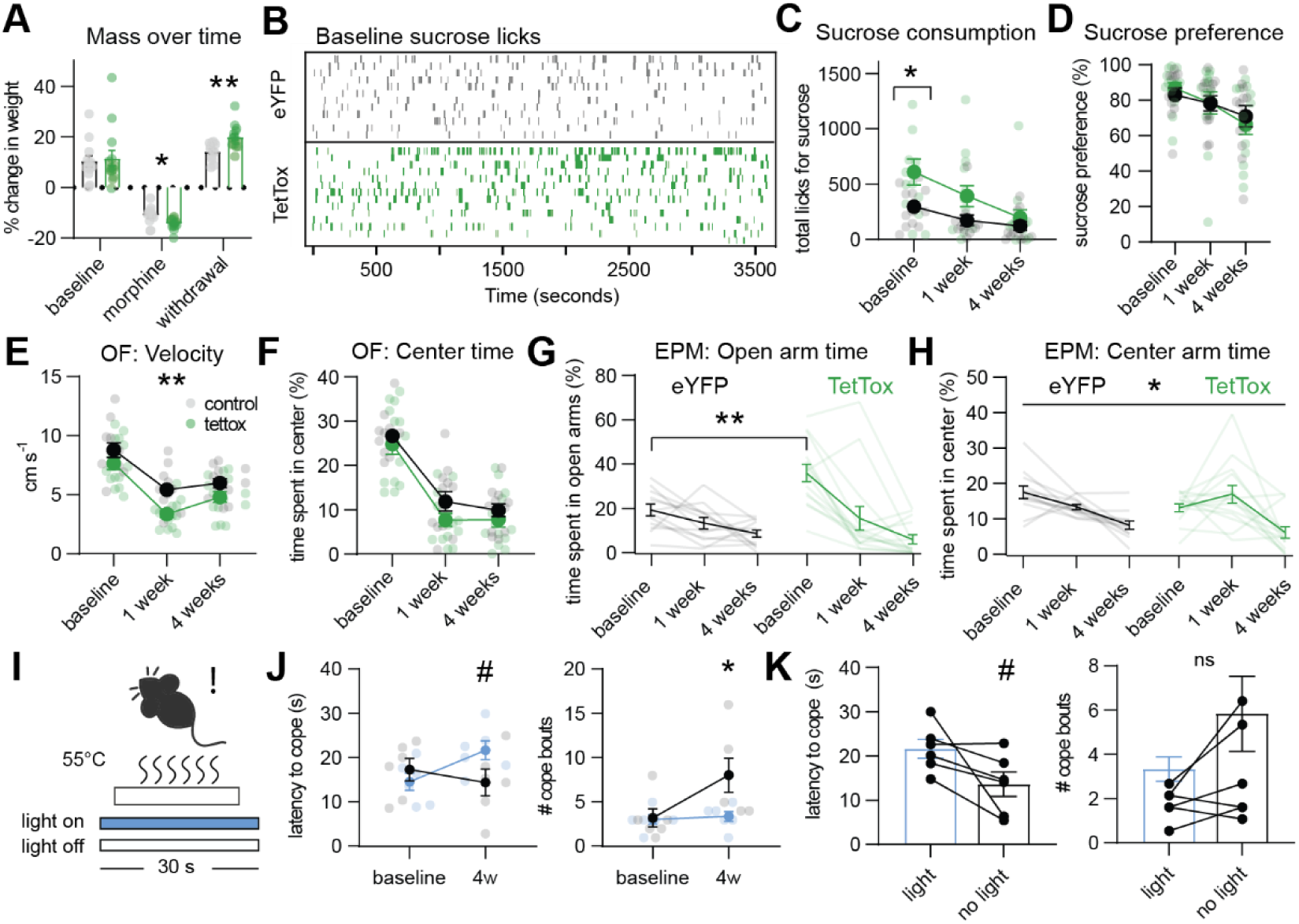
Additional TetTox behavioral tests. (A) Mass of mice throughout the experiment. Change in mass was calculated for different epochs of time. Baseline change was measured as the percent between between pre-surgery weights and ∼ 8 weeks later. Change during morphine escalation was measured as the percent change from the first day to the final day of injections (7 days). Finally, change during withdrawal was measured as the percent change from the first day of withdrawal to 30 days later. 2-way RM ANOVA followed by Šídák’s multiple comparisons; interaction: F_(2,48)_ = 3.252, p = 0.0474; control vs TetTox morphine, p = 0.0364; control vs TetTox withdrawal, p = 0.0059. (B) Sucrose consumption. Raster plot indicating licks performed by control and TetTox mice over a 1-hour period prior to morphine exposure. (C) Sucrose consumption. TetTox mice consume more sucrose than control mice at baseline, but there was no interaction between TetTox and withdrawal time (F_(2, 48)_ = 1.487, p = 0.2363). (D) Sucrose preference. There was no interaction between TetTox treatment and withdrawal time (2-way RM ANOVA with Šídák’s multiple comparisons: F_(2, 48)_ = 1.487, p = 0.2363). (E) Open field velocity. There is no interaction between TetTox and withdrawal time (F_(2,48)_ = 0.9216, p = 0.4048), but TetTox does influence velocity (F (2, 48) = 8.858, p = 0.0066) after 1 week of withdrawal (p = 0.0083). (F) Open field center time. There is no interaction between TetTox and withdrawal time (F_(2,48)_ = 0.2433, p = 0.7850) or main group effect (F_(1, 24_) = 3.678, p = 0.0671). (G) Elevated plus maze open arm time. There is an interaction between TetTox and withdrawal time (F_(2,48)_ = 6.409, p = 0.0034), such that TetTox elevates time spent in the open arms at baseline (p = 0.0019). (H) Elevated plus maze center time. 2-way RM ANOVA followed by Šídák’s multiple comparisons; interaction: F_(2,48)_ = 3.248, p = 0.0475. No significant post-hoc comparisons. (I) Hot plate assay. Mice were exposed to 55°C for 30 s during laser stimulation (20 hz, 5 ms pulse, 5-7 mW blue light). (J) Left, cope latency. 2-way RM ANOVA followed by Šídák’s multiple comparisons; interaction: F_(1,10)_ = 4.740, p = 0.0545; eYFP vs Chr2 at 4w, p = 0.0947. Right, number of cope bouts. 2-way RM ANOVA followed by Šídák’s multiple comparisons; group effect: F_(1,10)_ = 6.749, p = 0.0291; eYFP vs Chr2 at 4w, p = 0.0202. (K), Chr2 mice only, blue light stimulation vs no light. Left, paired t-test: t_(5)_ = 2.388, p = 0.0625. Right, paired t-test: t_(5)_ = 1.746, p = 0.1412.

**Figure S7.**
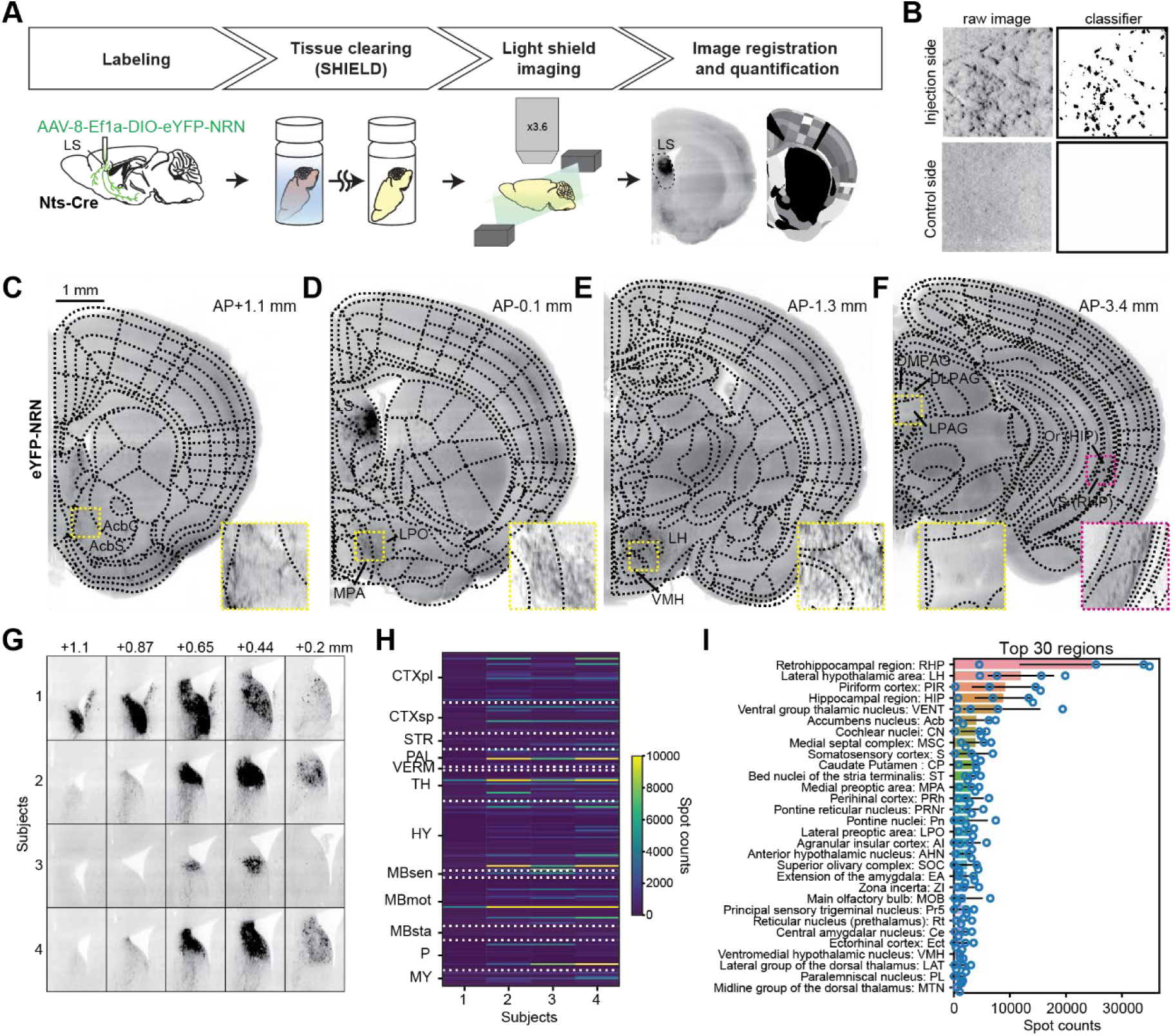
LS-Nts projection targets. (A) SHIELD Tissue clearing pipeline. In brief, 4 *Nts*-Cre mice (3 females, 1 male) were unilaterally injected with AAV8-EF1a-DIO-eYFP-NRN in the LS and allowed to recover for 3 weeks. Brains were removed and cleared using the SHIELD tissue clearing protocol, and cleared tissue was imaged using light sheet microscopy. Scans were registered to a common brain atlas and underwent classifier-based detection of synaptic puncta. (B) Example detection of both positive (top) and negative (bottom) signal. (C-F) Representative images of LS-*Nts* projections throughout the brain. (G-H) Histology and projection profile of all 4 subjects. (I) Quantification of spots (puncta) detected for the top 30 enriched brain regions.

